# Neuronal death in pneumococcal meningitis is triggered by pneumolysin and pilus-1 interactions with β-actin

**DOI:** 10.1101/2020.08.20.258681

**Authors:** Mahebali Tabusi, Sigrun Thorsdottir, Maria Lysandrou, Ana Rita Narciso, Melania Minoia, Birgitta Henriques-Normark, Federico Iovino

## Abstract

Neuronal damage is a major consequence of bacterial meningitis, but little is known about mechanisms that lead to neuronal death. *Streptococcus pneumoniae* (pneumococcus) is a leading cause of bacterial meningitis and many survivors develop neurological sequelae after the acute infection has resolved, possibly due to neuronal damage. Here, we studied mechanisms for pneumococcal interactions with neurons. Using human primary neurons and co-immunoprecipitation assays, we showed that pneumococci interact with the cytoskeleton protein β-actin through the pilus-1 adhesin RrgA and the cytotoxin pneumolysin (Ply), thereby promoting adhesion and uptake into neurons and neuronal death. Using our bacteremia-derived meningitis mouse model, we observed that RrgA- and Ply-expressing pneumococci co-localize with neuronal β-actin. We found that pneumococcal-infected neurons show increased intracellular Ca2+ levels depending on RrgA and mainly Ply which likely cause actin cytoskeleton disassembly leading to neuronal damage. Finally, neuronal death caused by pneumococcal infection could be inhibited using antibody against β-actin.

## Introduction

The Gram-positive bacterium *Streptococcus pneumoniae* (the pneumococcus) is the main cause of bacterial meningitis worldwide (***Iovino et al., 2016a***; ***Mook-Kanamori et al., 2011***). Despite access to antibiotics and intensive care, the mortality rate in pneumococcal meningitis remains high, ranging from 10-40% depending on geographical region (***van de Beek et al., 2006***; ***van de Beek et al., 2004***). Moreover, around 50% of the survivors suffer from permanent neurological damages, sequelae, after the infection has resolved (***Lucas et al., 2016***; ***Schmidt et al., 2016***; ***Chandran et al., 2011***; ***Nau et al., 2002***). Little is known on mechanisms for how pneumococci interact with and damage neurons.

Pneumococcal infections usually start with pneumococcal colonization of the upper respiratory tract. From this location bacteria may reach the blood stream and then interact and by-pass the blood-brain barrier (BBB) endothelium to cause meningitis (***Iovino et al., 2016a***; ***Iovino et al., 2013***). It has been shown that the RrgA protein of the pneumococcal pilus-1 interacts with the endothelial receptors pIgR and PECAM-1, thereby promoting pneumococcal spread through the BBB and infection of the brain (***Iovino et al., 2016b***). Pilus-1 has previously been shown to promote colonization, virulence, and pro-inflammatory responses in mouse models (***Barocchi et al., 2006***), and it is composed of three structural proteins, RrgA, RrgB and RrgC, where the tip pilin protein RrgA has been found to be the major adhesin to epithelial cells (***Nelson et al., 2007***), and RrgB to be the major stalk protein (***Barocchi et al., 2006***). The cytotoxin of pneumococci, pneumolysin (Ply), has also been shown to affect the brain. Indeed, purified Ply was observed to induce microglial and neuronal apoptosis (***Hirst et al., 2004***; ***Braun et al., 2002***). Ply is a cholesterol-dependent and pore-forming toxin that has been found to induce pro-inflammatory responses, but recently we showed that in certain cell types such as alveolar macrophages and dendritic cells, Ply may also induce anti-inflammatory responses and effect T-cell responses (***Subramanian et al., 2019***).

Pneumococci interact with host cells by adhering to their surfaces and by becoming internalized into certain cells such as immune cells. Like other bacteria they may exploit transport systems of host cells as mechanisms for adhesion and uptake. For example, both pneumococcal RrgA and the surface protein PspC have been shown to bind to pIgR (polymeric immunoglobulin receptor) on human nasopharyngeal epithelial and brain endothelial cells, and the physiological function of pIgR is to transport immunoglobulins across human-cell barriers (***Iovino et al., 2017***; ***Johansen et al., 2001***; ***Zhang et al., 2000***). In eukaryotic cells, the actin skeleton is essential for cellular development, metabolism, and immunity (***Krauss et al., 2005***), and β-actin is a predominant isoform among the actin cytoskeleton proteins (***Vedula et al., 2018***; ***Khaitlina et al., 2001***), essential for cell growth, migration, and the G-actin pool (***Bunnell et al., 2011***; ***Haglund et al., 2011***). In neurons, one of the main functions of actin filaments is the transport of microtubules along axons which is fundamental for neuronal development (***Hasaka et al., 2004***). Some bacterial pathogens, such as *Shigella* and enteropathogenic *Escherichia coli*, have been shown to exploit cytoskeletal proteins in eukaryotic cells to activate host-mediated uptake (***Croxen et al., 2010***; ***Yoshida et al., 2006***). However, little is known about pneumococcal interactions with the actin skeleton. It has been demonstrated though that Ply has direct transmembrane interactions with actin within the lipid bilayer of the plasma membrane of astrocytes (***Hupp et al., 2013***).

Here, we investigated pneumococcal interactions with neurons. We found that the pilus-1 protein RrgA can bind to neurons. Moreover, we show that presence of RrgA and the cytotoxin Ply promote pneumococcal entry into neurons. Also, we found that RrgA and Ply cause neuronal death, and that both proteins interact with β-actin of the neuronal cytoskeleton and induce Ca2+ release, thereby potentially damaging the cytoskeleton and the cells. We finally showed that the neurotoxic effect can be blocked using anti-β-actin antibody. The generated data potentially helps to explain mechanisms for why pneumococci frequently cause chronic neurological sequelae.

## Results

### Neuronal cell death by *S. pneumoniae* infection is pilus-1 and pneumolysin dependent

We first investigated the importance of pilus-1 and Ply in neuronal death. As *in vitro* model for neurons we used SH-SY5Y human neuroblastoma cells, and neurons obtained by differentiating SH-SY5Y cells using retinoic acid as previously described (***Shipley et al., 2016***). Validation of neuronal differentiation was assessed by detection of neuronal markers MAP2 and NSE (***Soltani et al., 2005***; ***Isgró et al., 2015***) by western blot and immunofluorescence staining (***Supplementary Figures 1A*** and **1B**) and by morphological microscopy analysis (***Supplementary Figure 1C***). Neurons were challenged with the pneumococcal strain TIGR4 and then stained with a live/dead dye and visualized using high-resolution live-cell imaging microscopy. We observed that the neurons were rapidly killed by the TIGR4 strain. Thus, 2 hours post-infection all imaged neurons showed clear signs of cell death (***Figure 1A***). To study the roles of pilus-1 and Ply, we used mutant strains lacking the pilus-1 islet, TIGR4Δ*rrgA*-*srtD*, (***Iovino et al., 2016***) or Ply, TIGR4Δ*ply* (***Subramanian et al., 2019***). When infected with either of the two mutants, neurons showed a significant decrease in cell death compared with wild-type (wt) TIGR4 infection (***Figures 1A*** and ***1B***). Furthermore, neuronal cell death caused by TIGR4Δ*ply* was approximately 50% lower than what was observed for TIGR4Δ*rrgA*-*srtD* (***Figure 1C***). These data suggest that the neuronal damage caused by Ply, which could act as toxin upon being released after bacterial lysis or while being attached on the pneumococcal surface (***Subramanian et al., 2019***; ***Braun et al., 2002***), is more severe than the damage caused by pilus-1, which mediate physical interaction with the host cell. After 2 hours infection with the double mutant TIGR4Δ*rrgA*-*srtD*Δ*ply*, the neuronal death rate was less than 5% of wt TIGR4. and considerably lower than infection with either of the single mutants TIGR4Δ*rrgA* or TIGR4Δ*ply* (***Figures 1A, 1B*** and ***1C***). Thus, we conclude that both pilus-1 and Ply contribute to neuronal cell damage.

**Figure 1.**
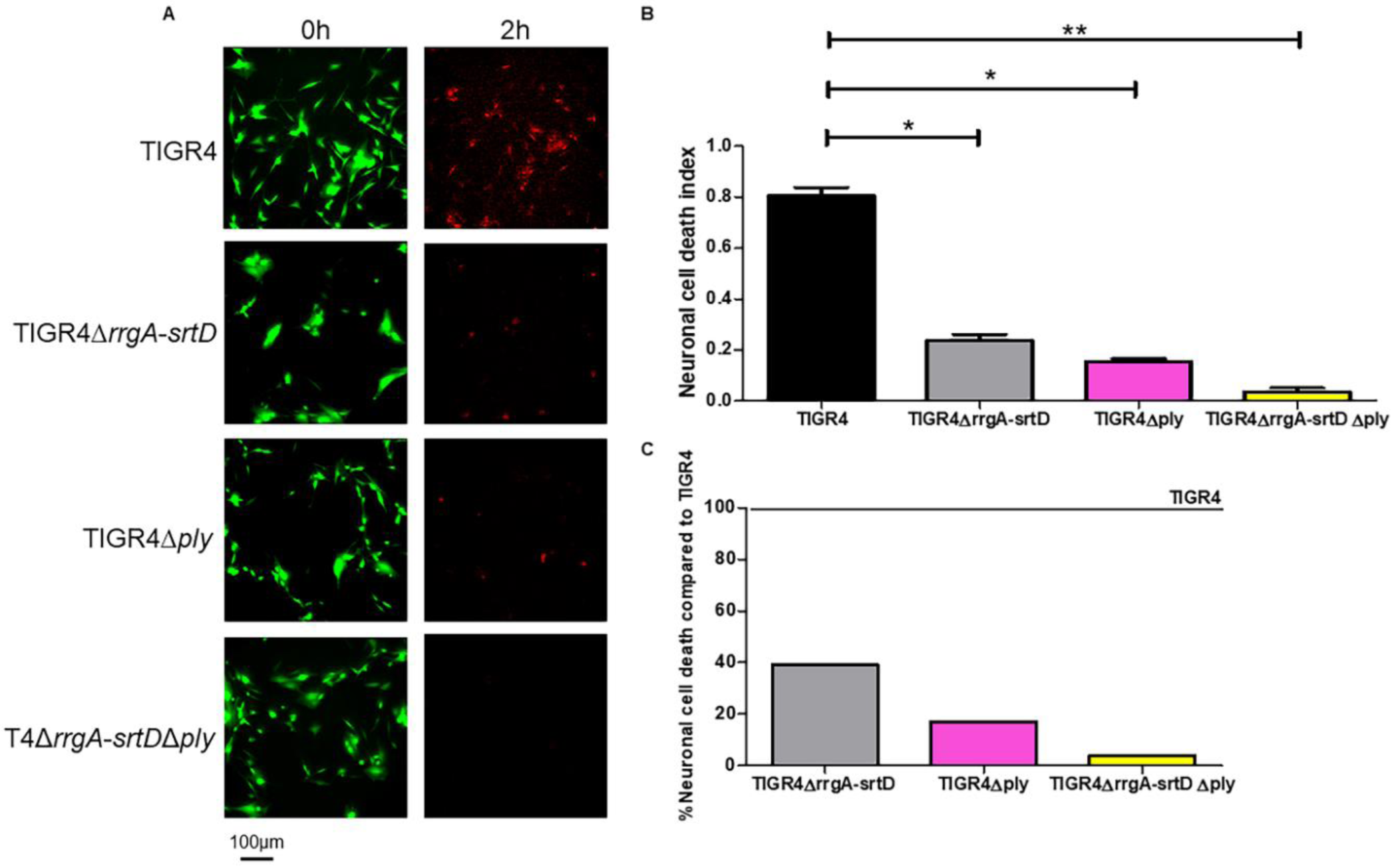
Presence of the pneumococcal pilus-1 and Ply enhance neuronal cytotoxicity. (**A**) Images from live-cell imaging at the start (time 0) and at the end, 2 hours post-infection. Differentiated neurons stained with LIVE/DEAD dye express green fluorescence when they are alive, and a red fluorescence when undergoing cell death. (**B**) Quantification of the neuronal cell death in the 2-hour infection experiment shown in Figure 1A; Green (488 nm) / Red (594 nm) represents the neuronal cell death index, calculated by dividing the total area occupied by the green fluorescence signal at time 0 by the total area occupied by the red fluorescence signal at the end of the infection. Per each pneumococcal strain, a total of 2 biological replicates (2 wells with neurons, each well seeded in a different day) have been used for the 1-hour experiment, and a total of 2 biological replicates (2 wells with neuron, each well seeded in a different day). Columns in the graphs represent average values, error bars represent standard deviations. ** = p<0.001, * = p<0.05. (**C**) Percentage average values of neuronal cell death calculated setting the average value of neuronal cell death of TIGR4 to 100%. The percentage average values were calculated using the neuronal cell death index values shown in Figure 1B.

### The pilus-1 adhesin RrgA mediates binding to and promotes internalization by neurons while Ply only enhances uptake

Next, we studied whether pilus-1 and Ply influence pneumococcal adhesion to and internalization by neurons. First, we infected SH-SY5Y cells or neurons with wt TIGR4 and found that pneumococci can adhere to both differentiated neurons and SH-SY5Y before differentiation (***Figure 2A*** and ***Supplementary Figure 2A***). Then we used our pilus-1 deletion mutant, TIGR4Δ*rrgA*-*srtD*, and observed that it adhered significantly less to neurons and SH-SY5Y cells as compared to wt TIGR4 (***Figure 2A*** and ***Supplementary Figure 2A***), suggesting that pilus-1 promotes pneumococcal interaction with neurons. High-resolution immunofluorescence microscopy analysis confirmed that higher numbers of TIGR4 bacteria bound to neurons and SH-SY5Y cells than the non-piliated TIGR4Δ*rrgA*-*srtD* (***Figures 3A-B*** and ***Supplementary Figure 3A-B***). To study if the pilin adhesin RrgA influences the adhesion to neurons, we used a mutant lacking the tip protein RrgA, TIGR4Δ*rrgA* (***Iovino et al., 2016***). We found that adhesion to neurons was significantly reduced as compared to wt TIGR4 (***Figure 2A*** and ***Supplementary Figure 2A***), and reached similar levels as using the pilus-1 mutant TIGR4Δ*rrgA*-*srtD* (***Figure 2A*** and ***Supplementary Figure 2A***). Bacterial adherence is a prerequisite for subsequent internalization by host cells. Internalization assays using both neurons and undifferentiated SH-SY5Y cells showed that the mutants TIGR4Δ*rrgA*-*srtD*, and TIGR4Δ*rrgA*, but not the complemented strain TIGR4Δ*rrgA+RrgA* (***Nelson et al., 2007***), were taken up significantly less than wt TIGR4 (***Figure 2B*** and ***Supplementary Figure 2B***). Thus, RrgA is the structural component of pilus-1 that mediates pneumococcal interaction with neurons.

**Figure 2.**
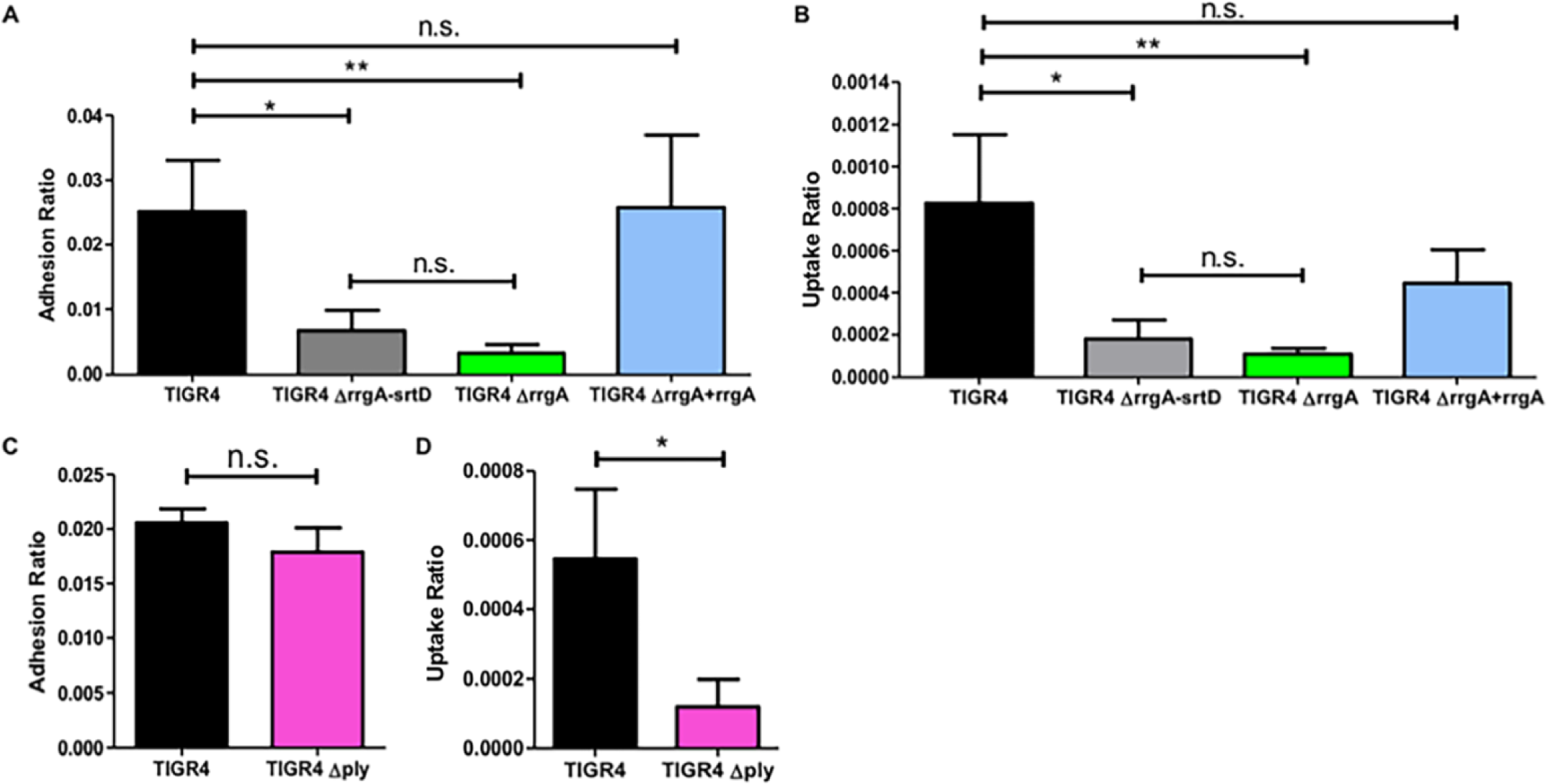
RrgA promotes pneumococcal binding to neurons, and both RrgA and Ply increases pneumococcal uptake into neuronal cells. Pneumococcal interactions with neurons was investigated *in vitro* using differentiated neurons and infection with (**A and B**) wt TIGR4 and its isogenic mutants, TIGR4Δ*rrgA*-*srtD*, TIGR4Δ*rrgA*, and TIGR4Δ*rrgA*+*rrgA*, and in (**C and D**) wt TIGR4 and TIGR4Δ*ply*. (A) CFU-based adhesion to neurons. (B) Uptake into neurons. (C) Adhesion ratio was calculated as [CFU of adhered bacteria] / [CFU of (non−adhered bacteria + adhered bacteria)]. (D) Uptake ratio was calculated as [CFU of invaded bacteria] / [CFU of adhered bacteria)]. For all graphs (A-D) the columns represent average values, and error bars represent standard deviations. Each graph shows an overview at least three (n≥3) biological replicates. ** = p<0.001, * = p<0.05, n.s. = non-significant.

**Figure 3.**
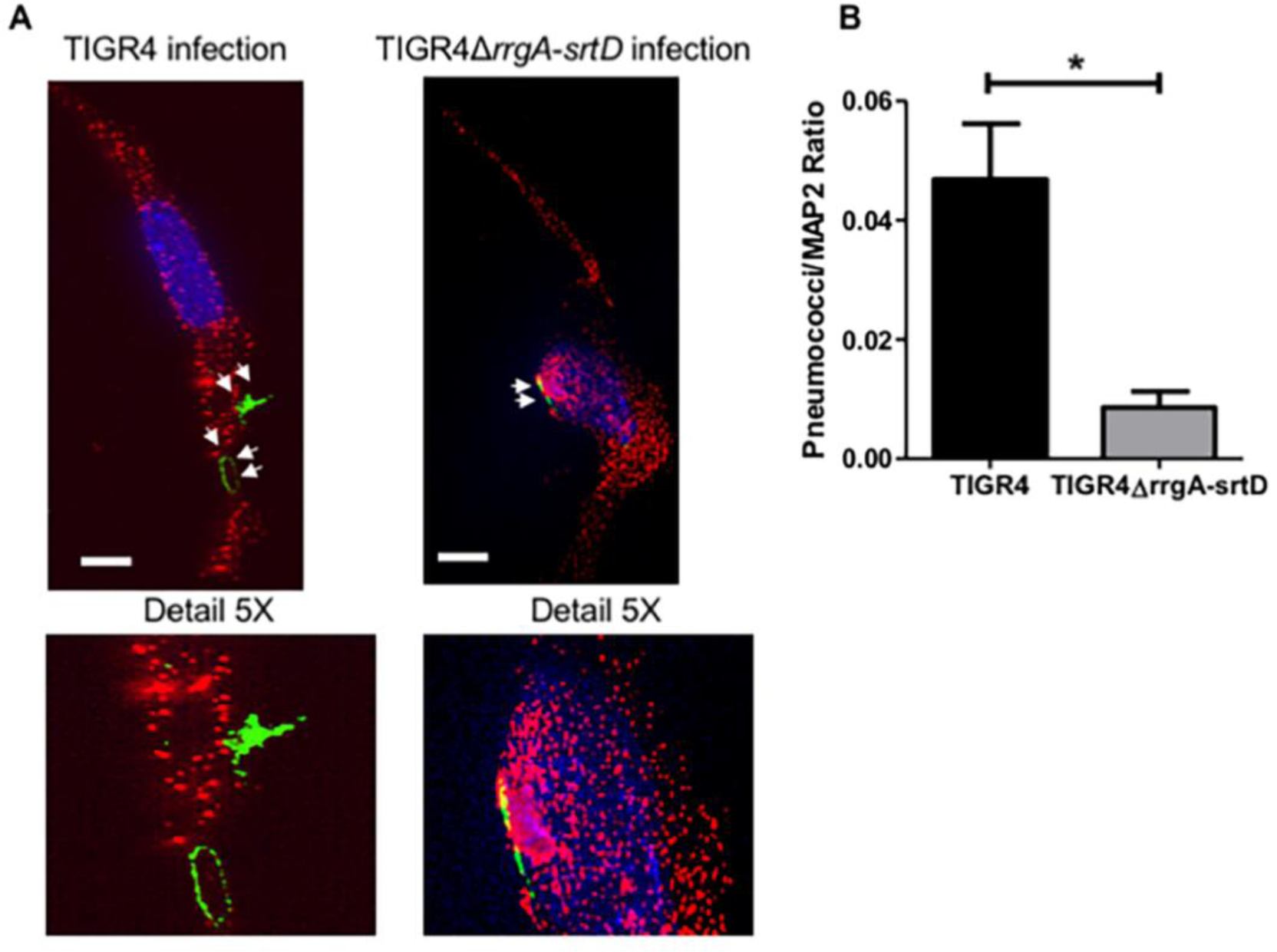
High-resolution fluorescence microscopy analysis supports that pilus-1 expression increases pneumococcal adhesion to neurons. (**A**) Piliated TIGR4 bacteria or its isogenic mutant in pilus-1 TIGR4Δ*rrgA*-*srtD* were used to infect neuronal cells. High-resolution fluorescence microscopy analysis was performed on adhered bacteria where neurons were stained with anti-MAP2 antibody combined with goat anti mouse Alexa Fluor 594 (red), and the pneumococcal capsule stained with anti-serotype 4 capsule antibody combined with goat anti rabbit Alexa Fluor 488 (green). White arrows point to pneumococci that adhered to neurons. White scale bars represent 10 µm. Two representative images are shown selected from 200 cells with adhered bacteria imaged per pneumococcal strain. The panel “Detail 5X” displays a 5X-magnified image of the area in the original images where the bacteria adhered to neurons. (**B**) Quantification analysis of the amount of pneumococcal signal detected on the plasma membrane of neurons by high-resolution microscopy images shown in Figure 3A. For quantification of the bacterial fluorescence signal on neurons, in each image (n=200 neurons with adhered bacteria per each pneumococcal strain) the area occupied by the green fluorescence signal of the bacteria was divided by the area occupied by the red fluorescence signal of neurons. All areas were measured in square pixels and calculated with the software Image J. Columns in the graph represent average values. Error bars represent standard deviations. The Pneumococci/MAP2 ratio is shown on the Y axis; * = p<0.05.

In contrast, the mutant strain lacking Ply, TIGR4Δ*ply*, adhered to neurons and SH-SY5Y cells to a similar extent as wt TIGR4 (***Figure 2C*** and ***Supplementary Figure 2C***). However, absence of Ply expression resulted in a significantly lower uptake by the neurons and SH-SY5Y cells(***Figure 2D*** and ***Supplementary Figure 2D***), suggesting that Ply, through its pore-forming action on the neuronal plasma membrane, can promote bacterial uptake into neurons.

### Both RrgA and Ply interact with neuronal β-actin

To identify proteins on the plasma membrane of neurons that bind to RrgA and/or Ply, we setup a co-immunoprecipitation assay using His-tagged RrgA and Ply coupled to Ni-NTA magnetic beads incubated with a cell lysate of differentiated human neurons. The quality of the neuronal cell lysate was assessed by SDS-page and Coomassie staining (***Supplementary Figure 4***). We found that the cytoskeleton protein β-actin was the only abundant neuronal protein that bound to RrgA and Ply that was not present in the negative control (***Figure 4A*** and ***Supplementary Tables 1-3***). α tubulin proteins also bound to RrgA and Ply at high scores, but all of them were present in the negative control (***Supplementary Table 3*** and ***Supplementary Figure 5***). To confirm the specificity of the β-actin interaction, we performed co-immunoprecipitation experiments using recombinant β-actin protein. Western blot analysis showed that both recombinant RrgA and Ply, but not the pilus-1 backbone protein RrgB, bound to β-actin (***Figure 4B***). To assess whether RrgA and Ply were interacting exclusively with β-actin and not with other cytoskeleton proteins, we made use of the α-tubulin-1B chain. Co-immunoprecipitation experiment showed that neither RrgA nor Ply bind to α-tubulin-1B, as only a faint α-tubulin-1B band was detected that also appeared in the control with only Ni-NTA beads, indicating an unspecific affinity of α-tubulin-1B for the beads (***Figure 4B***). Taken together our results suggest that both RrgA and Ply bind specifically to neuronal β-actin.

**Figure 4.**
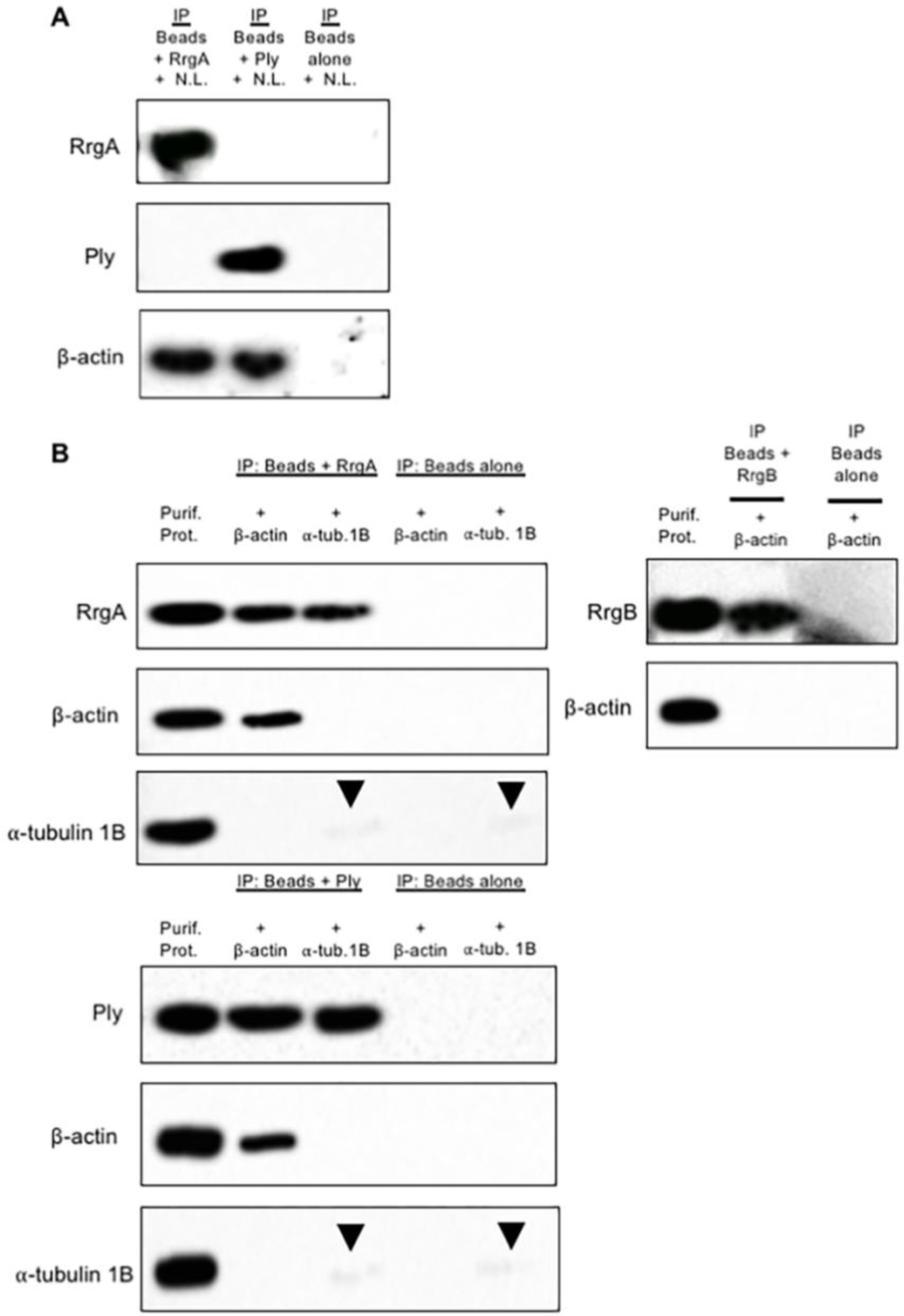
Co-immunoprecipitation shows that RrgA and Ply interact with β-actin of neurons. (**A**) Western blot analysis performed using the protein content obtained after the co-immunoprecipitation (IP) experiment using Ni-NTA beads coupled with purified RrgA or Ply and incubated with neuronal lysate (N.L.). Three separate western blots using the same samples were performed for the detection of RrgA, Ply and β-actin respectively. (**B**) Western blot analysis performed after co-immunoprecipitation experiments using purified RrgA and Ply incubated with the purified recombinant proteins β-actin or α-tubulin 1B (as negative control). Black arrows point to faint bands that were detected for α-tubulin 1B. The presence of these bands of similar intensity in the IP samples with beads coupled with RrgA or Ply and beads alone indicates a slight unspecific affinity of α-tubulin 1B for the Ni-NTA beads. Four separate western blots using the same samples were performed for detection of RrgA, Ply, β-actin and α-tubulin 1B. As a specificity control, western blot analysis was also performed after the co-immunoprecipitation experiments using purified RrgB incubated with the purified recombinant proteins β-actin. Two separate western blots using the same samples were performed for detection of RrgB and β-actin.

**Figure 5.**
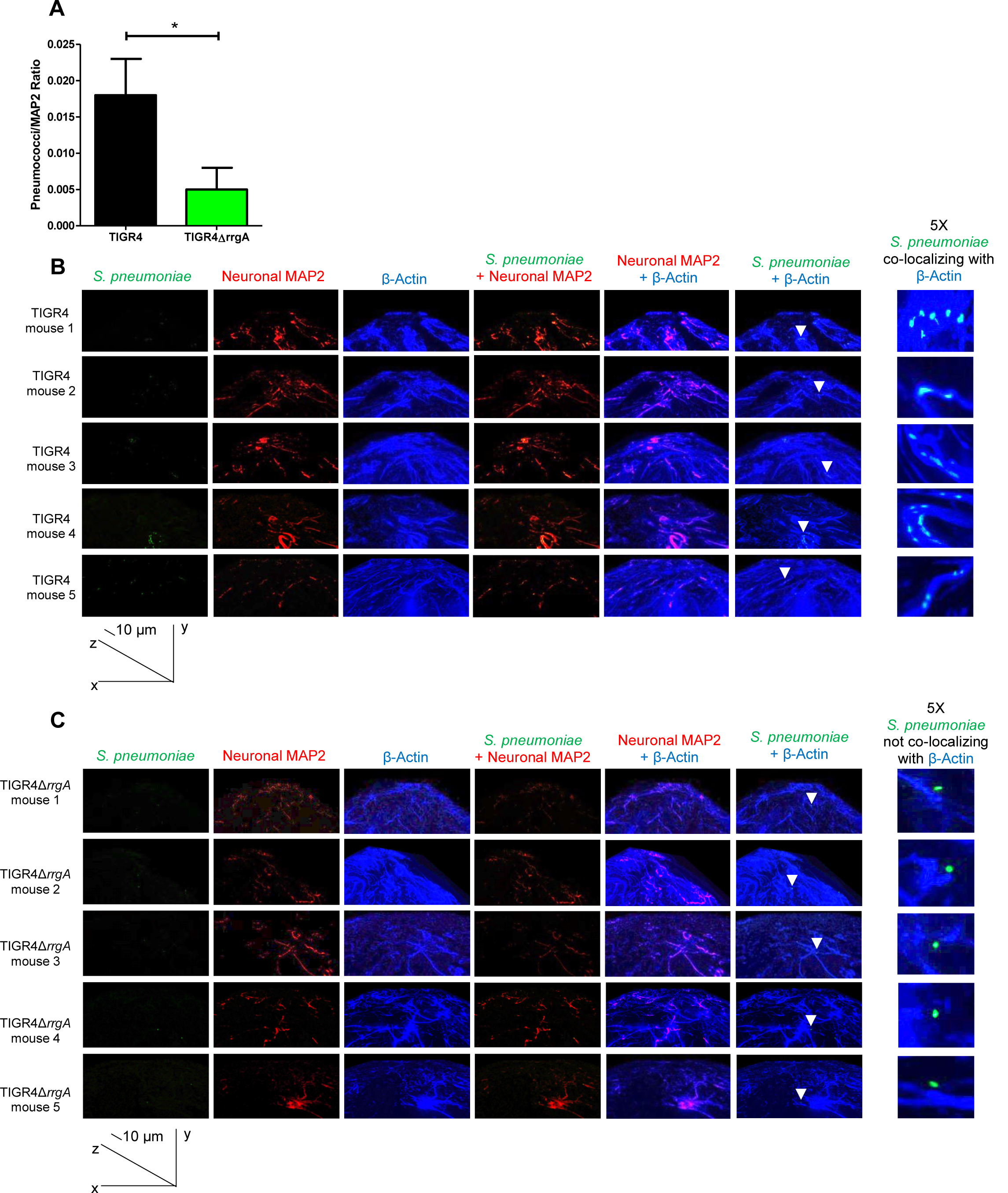
Mouse brain tissue *ex vivo* analyzed by high-resolution fluorescence microscopy and 3D reconstruction imaging showing that pneumococci associated with neurons co-localize with neuronal β-actin only when expressing RrgA. Pneumococci were stained with anti-serotype 4 capsule antibody combined with goat anti rabbit Alexa Fluor 488 (green), neurons were stained with anti-MAP2 antibody labelled with Zenon labeling mouse IgG 594 fluorophore (red), β-actin was stained with anti-β-actin antibody combined with goat anti mouse Alexa Fluor 647 (far red, a blue color was assigned using the Softworx imaging software); high-resolution microscopy analysis and 3D reconstruction imaging (Volume viewer function of the Softworx imaging software) was performed to firstly detect pneumococci in the brain tissue of the mice co-localized with neurons, then to analyze the co-localization between pneumococci and β-actin, and analysis of β-actin with the neuronal marker MAP-2 to distinguish β-actin of neurons co-localization. (A) For quantifying the bacterial fluorescence signal on neurons, in each image the area occupied by the green fluorescence signal of the bacteria was divided by the area occupied by the red fluorescence signal of neurons, all areas were measured in square pixels and calculated with the software Image J; columns in the graph represent average values, error bars represent standard deviations, the Pneumococci/MAP2 ratio is shown on the Y axis; * = p<0.05. (B and C) At the bottom left corner of figures in panels B and C the graph shows the angle of the 3D reconstruction of each image (XYZ axes) and the scale bars; six tissue sections for each mouse (5 mice) infected with either TIGR4 (A) or TIGR4Δ*rrgA* (B) were imaged, and per each section twenty-five images in random regions of the section were taken. White arrows in the panel “S. pneumoniae + β-actin” point towards specific area of the tissue sections to highlight the co-localization between piliated pneumococci and β-actin (A), and the absence of co-localization between non-piliated pneumococci and β-actin (B); the panel “5X S. pneumoniae co-localizing with β-actin” displays the region of brain tissue in close proximity of the white arrows with an enhanced 5X magnification.

### Pneumococci expressing RrgA co-localize with β-actin of neurons *ex vivo*

To investigate if RrgA mediates pneumococcal binding to neurons *in vivo*, we made use of our bacteremia-derived meningitis mouse model (***Iovino et al, 2016a***; ***Iovino et al., 2016b***; ***Iovino et al, 2017***). Brain tissue sections from mice infected with TIGR4 or its isogenic mutant TIGR4Δ*rrgA* were examined by high-resolution immunofluorescence microscopy and 3D imaging reconstruction. A significantly stronger bacterial fluorescence signal associated with neurons was observed in the brain of mice infected with wt TIGR4 as compared to mice infected with TIGR4Δ*rrgA* (***Figure 5A***). Furthermore, we found that TIGR4 co-localized with β-actin of neuronal cells, stained using the specific marker MAP2 (***Figure 5B***), while infection with the mutant TIGR4Δ*rrgA* showed no co-localization between the bacteria and neuronal β-actin. Hence, these *in vivo* data support our *in vitro* data, suggesting that pneumococcal RrgA interacts with β-actin on neurons.

### Pneumococci exploit β-actin filaments to become internalized by neurons

To study internalization into neurons, we used a CFU-based invasion assay. We found that the double mutant TIGR4Δ*rrgA-srtD*Δ*ply*, in contrast to wt TIGR4, was poorly internalized by neurons (***Figure 6A***). Infected neurons with intracellular pneumococci were fixed, permeabilized and immunofluorescence staining was performed to detect *S. pneumoniae* cells and β-actin. Quantification analyses confirmed that TIGR4Δ*rrgA-srtD*Δ*ply* bacteria were found only rarely inside neurons (***Figure 6B***). High-resolution fluorescence microscopy combined with orthogonal view analysis clearly demonstrated that intracellular TIGR4 pneumococci co-localized with β-actin staining (***Figure 6C***), while the very few intracellular TIGR4Δ*rrgA-srtD*Δ*ply* bacteria observed did not colocalize with neuronal β-actin (***Figure 6D***). These results argue that pneumococci, once bound to β-actin on the plasma membrane, retain this binding after becoming internalized, and suggest that pneumococci exploit the cytoskeleton β-actin filaments to enter neurons. The β-actin staining in close proximity to intracellular bacteria was much more intense than the β-actin staining in the rest of the neuronal cell (***Figure 6E***), suggesting that pneumococcal binding to β-actin on the neuronal plasma membrane could trigger actin polymerization, promoting pathogen internalization.

**Figure 6.**
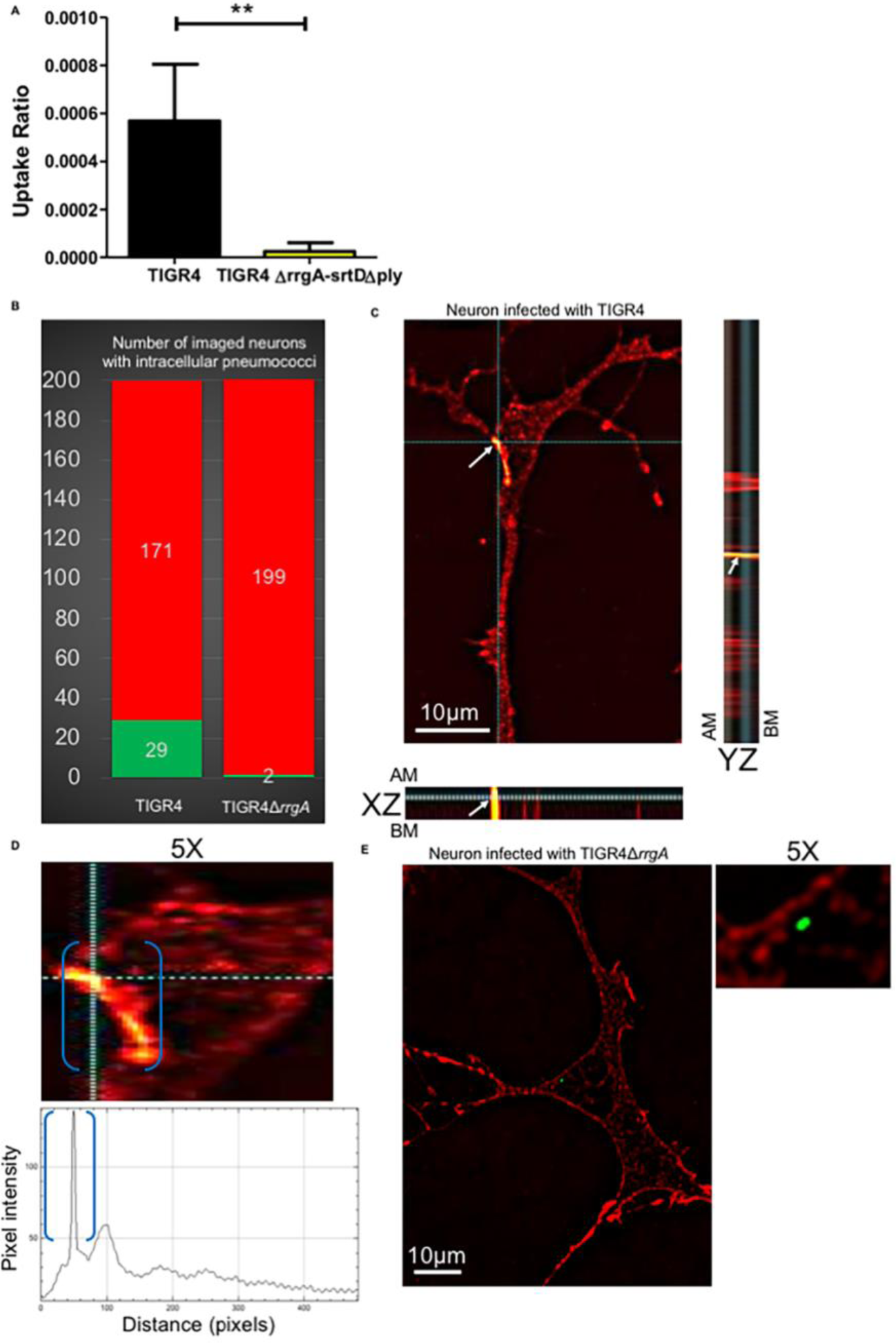
RrgA and Ply increase pneumococcal uptake by neurons and intracellular piliated pneumococci co-localize with neuronal β-actin. (**A**) CFU-based internalization assay using wt TIGR4 and its isogenic double mutant TIGR4Δ*rrgA*-*srtD*Δ*ply*. The uptake ratio by neurons was calculated as [CFU of intracellular bacteria] / [CFU of adhered bacteria)]. In the graph, columns represent average values, and error bars represent standard deviations. The graph shows an overview of three biological replicates. ** = p<0.001. (**B**) High-resolution fluorescence microscopy analysis was performed of neurons with intracellular pneumococci that were fixed, and permeabilized. Immunofluorescence staining was performed to detect β-actin with an anti-β-actin antibody combined with goat anti mouse Alexa Fluor 594 (red) and pneumococci with anti-serotype 4 capsule antibody combined with goat anti rabbit 488 (green). The graph shows a quantification of the number of neurons with intracellular pneumococci among a total number of 200 random neurons imaged per strain, either TIGR4 or TIGR4Δ*rrgA*. The green column represents the number of neurons with intracellular pneumococci and the red column the number of neurons without intracellular pneumococci. (**C**) Neurons with intracellular pneumococci after TIGR4 infection were imaged with z-stacks to capture the thickness of the neuronal cell (number z-stacks = 22). Intracellular pneumococci with the z-stack number = 9 was displayed from top view and in XZ-axes- and YZ-axes-orthogonal views to demonstrate intracellular localization of pneumococci (green). The imaged bacteria were within the neuronal cell thickness between the AM (apical membrane) and BM (basolateral membrane). Both XZ and YZ-axes-orthogonal views showed co-localization between pneumococci (white arrows) and intracellular β-actin. The image shown is a representative of 200 neurons imaged after TIGR4 infection. (**D**) 5X magnification of the neuron shown in Figure 6B focusing on the cell area in close proximity to intracellular pneumococci. The function Profile Plot of Image J was used to measure the intensity (pixels) of the red fluorescence signal of β-actin. Within blue brackets the β-actin staining in close proximity to intracellular pneumococci that corresponds to the pick of fluorescence intensity in the graph underneath the microscopy image is shown. (**E**) Neurons with intracellular pneumococci after TIGR4Δ*rrgA* infection were imaged with z-stacks to capture the thickness of the neuronal cell (number z-stacks = 22). The displayed image shows the z-stack number = 10. The panel “5X” shows the same image 5X magnified focusing on the area of neuronal cell in close proximity to the intracellular bacteria to highlight the absence of co-localization between TIGR4Δ*rrgA* and intracellular β-actin. This image is a representative of 200 neurons imaged after TIGR4Δ*rrgA* infection.

### Pneumococcal expression of Ply dramatically enhances intracellular calcium (Ca^2+^) levels of neurons, suggesting disruption of β-actin filaments

The interplay between the actin cytoskeleton and Ca^2+^ signaling was previously shown to play an important role during actin polymerization and neuronal growth and motility (***Cristofanilli and Akopian, 2006***; ***Gasperini et al., 2017***). Moreover, high levels of intracellular Ca^2+^ were described to be cytotoxic in neurons (***Stueber et al., 2017***; ***Kass and Orrenius, 1999***; ***Murphy et al., 1988***). Therefore, we performed Ca^2+^ imaging of neurons infected with wt TIGR4, the double mutant TIGR4Δ*rrgA-srtD*Δ*ply*, the single mutants TIGR4Δ*rrgA* and TIGR4Δ*ply*, and non-infected neurons, using the fluorescent Ca^2+^ indicator Fluo-8. We observed a significant increase of the Ca^2+^ flux level in neurons infected with wt TIGR4 compared to neurons infected with the double mutant TIGR4Δ*rrgA-srtD*Δ*ply* or non-infected neurons (***Figures 7A-C*** and ***7G***). While the level of intracellular Ca^2+^ release in neurons infected by TIGR4Δ*rrgA* was only moderately lower (not significant) than with TIGR4, absence of Ply caused a significant decrease of intracellular Ca^2+^ influx (***Figures 7D, 7E*** and ***7G***). Furthermore, when neurons were infected with TIGR4Δ*ply* in combination with purified Ply, the levels of intracellular Ca^2+^ release reached similar levels to that observed in neurons infected with wt TIGR4 (***Figure 7F***), suggesting that the increased Ca^2+^ influx is mainly caused by Ply released by pneumococci. An intact actin cytoskeleton was previously described to inhibit the activation of Ca^2+^ entry (***Rosado and Sage, 2000***), here we observed that, relative to TIGR4, the average values of intracellular Ca^2+^ peak intensities showed a 50% and 10-15% reduction for neurons infected with TIGR4Δ*ply*, and TIGR4Δ*rrgA* respectively (***Figure 7H***). Taken together these results suggest that RrgA and, in particular, Ply enhance the intracellular Ca^2+^ levels which possibly cause disruption of β-actin filaments finally leading to neuronal cell death.

**Figure 7.**
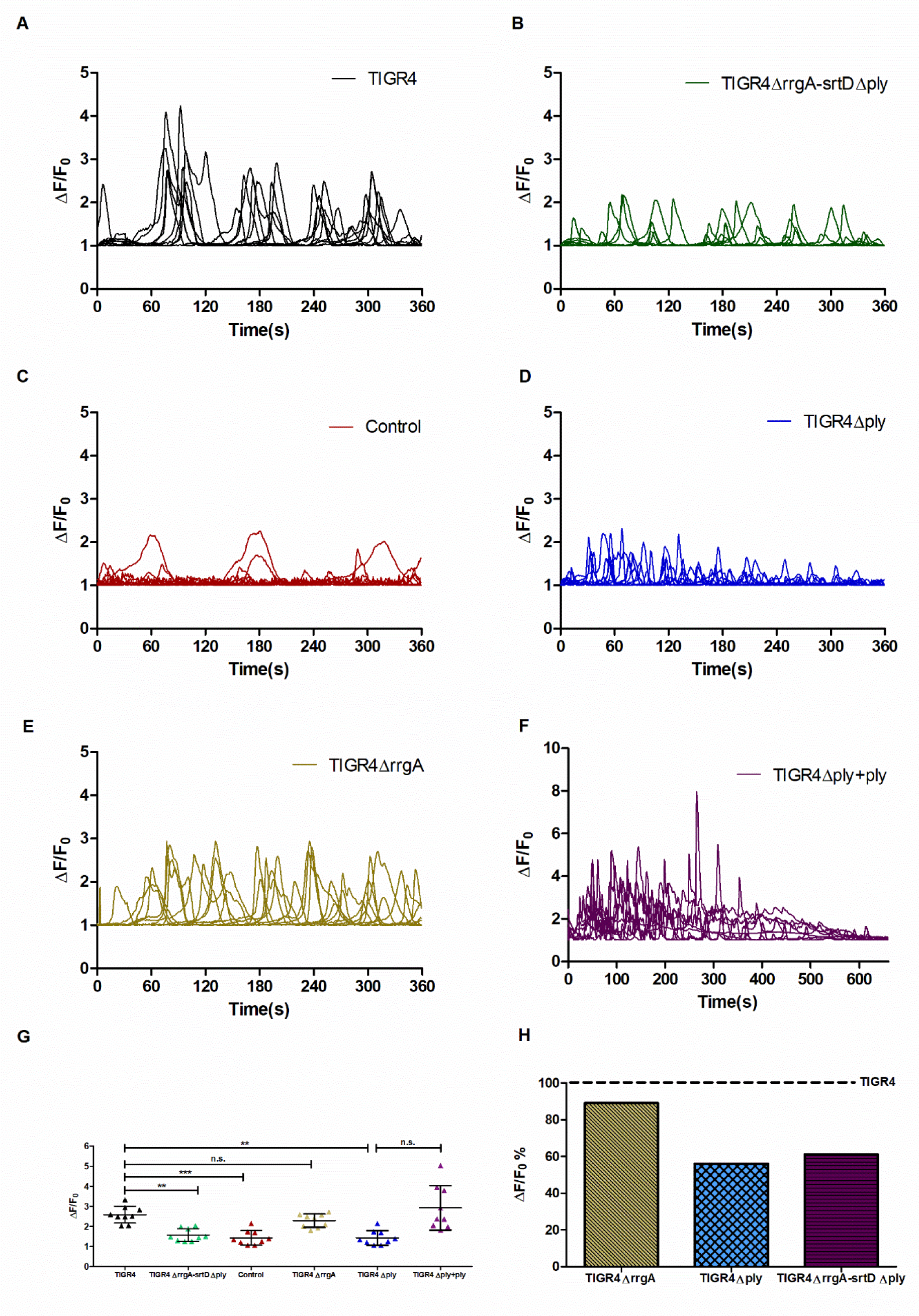
Neurons infected with Ply-expressing pneumococci showed increased levels of intracellular Ca^2+^. Intracellular Ca^2+^ release and influx in neurons infected with (**A**) TIGR4, (**B**)TIGR4Δ*rrgA*-*srtD*Δ*ply*, (**C**) non-infected neurons (Control), (**D**) TIGR4Δ*ply* and (**E**) TIGR4Δ*rrgA* were determined using Fluo-8 AM two hours post infection. Ca^2+^ imaging was performed for 6 minutes of time lapse in every 1 second using a FITC fluorescence channel. (**F**) Neurons were infected with TIGR4Δ*ply*, then 100ng/mL of Ply was added 2 min prior to imaging and Ca^2+^ imaging was performed for 11 minutes of time lapse in every 1 second using FITC fluorescence channel. In A-F, the Fluorescence intensity was measured using Image J from the region of interest (ROI) where a minimum of 10 ROI in each field was selected manually. ΔF/F0 was calculated from Image J-generated data (ΔF=Ft-F0, Ft represents the fluorescent intensity in each given time point and F0 represents the average fluorescent intensity of the resting value). Each graph displays the Ca^2+^ flux from 9 neuronal cells of 3 different experiments shown in 360 and 660 seconds. (**G**) Average ΔF/F0 values of relative peaks (n=9) during 6 minutes within ROI were shown to compare intracellular Ca^2+^ levels; ***= p<0.0001, **= p<0.001 n.s. = not-significant. Each data point in the graph represents one peak. (**H**) Percentage of average ΔF/F0 values of TIGR4Δ*rrgA*, TIGR4Δ*ply* and TIGR4Δ*rrgA*Δ*ply* compared to the ΔF/F0 values showed by neurons infected with wt TIGR4 (set to 100%). Each average value was calculated using the Ca^2+^ flux intensity values shown in Figure 7G (each dot in every group is one intensity value).

### Anti-β-actin antibody inhibits pneumococcal uptake and cellular toxicity in neurons

Anti-β-actin antibody was added to seeded neurons to block the bacterial binding site of β-actin on the neuronal plasma membrane. One-hour treatment with anti-β-actin antibody led to a significant reduction in the adhesion of TIGR4 bacteria to neurons, while adhesion after isotype control treatment was similar as what was observed in neurons without any antibody treatment (***Figure 8A***). Furthermore, TIGR4 adhesion was not affected by neuronal treatment with an antibody targeting the anaplastic lymphoma kinase (ALK), a receptor tyrosine kinase expressed on the neuronal plasma membrane (***Souttou et al, 2001***) (***Figure 8B***), indicating that the anti-β-actin antibody directly interferes with β-actin binding to pneumococcal RrgA.

**Figure 8.**
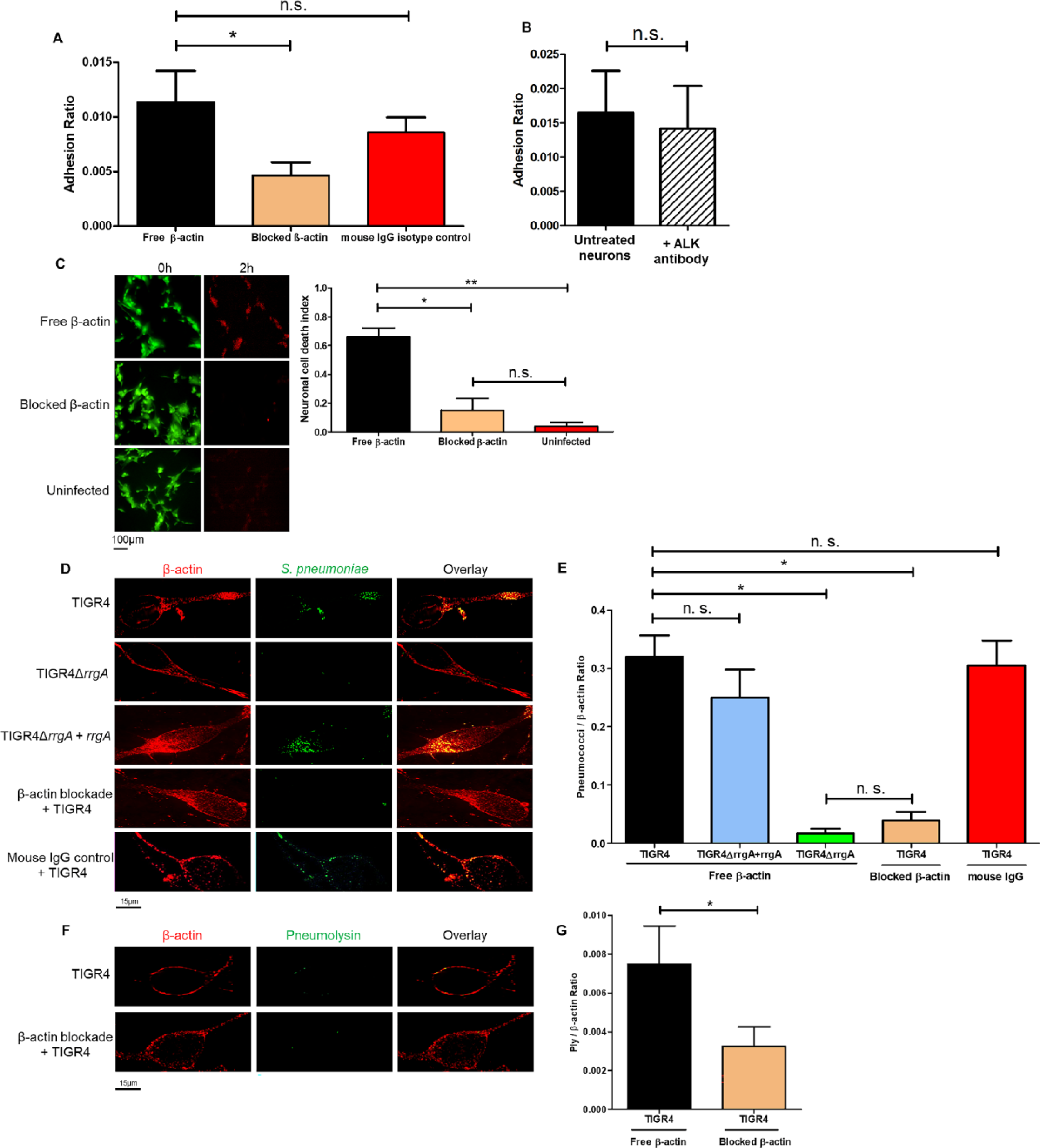
Blockade of β-actin on the plasma membrane with antibodies inhibit interactions with the cytotoxin Ply. CFU-based adhesion assays were performed using wt TIGR4 bacteria and in (**A**) untreated neurons, anti-β-actin-antibody-treated neurons and mouse-IgG-treated neurons, and in (**B**) untreated neurons and anti-ALK-antibody-treated neurons. The adhesion ratio was calculated as [CFU of adhered bacteria] / [CFU of (non−adhered bacteria + adhered bacteria)]. The columns represent average values, and error bars represent standard deviations. The graph shows an overview at three biological replicates. * = p<0.05, n.s. = non-significant. (**C**) Images from the live-cell imaging experiment at the start (0 h) and at the end, 2 hours post infection, and related quantification of neuronal cell death. Differentiated neurons were stained with a live/dead dye expressing green fluorescence for live cells and red fluorescence when undergoing cell death. Ratio Green (488 nm) / Red (594 nm) represents the neuronal cell death index, calculated by dividing the total area occupied by the green fluorescence signal at time 0 by the total area occupied by the red fluorescence signal at 2 hrs (as performed for data shown in Figure 1). Per each condition with pneumococci, a total of four biological replicates were used, and a total of two biological replicates were used for uninfected neurons. Columns in the graphs represent average values, and error bars represents standard deviations. ** = p<0.001, * = p<0.05, n.s= non-significant. (**D and F**) Infected neurons were fixed and stained with anti-β-actin antibody combined with goat anti mouse Alexa Fluor 594 for the detection of β-actin, and with anti-serotype 4 capsule antibody combined with goat anti rabbit Alexa Fluor 488 (green) for the detection of pneumococci (D). Anti-Ply antibody combined with goat anti rabbit Alexa Fluor 488 (green) was used for detection of Ply (F). Each neuron imaged in D and F is representative of 200 neurons imaged per each pneumococcal strain. (**E and G**) Quantification analysis of the amount of pneumococcal signal detected on the plasma membrane of neurons by high-resolution microscopy. For quantification of the (E) pneumococcal or (G) Ply fluorescence signal on neurons, in each image (n=200 neurons with adhered bacteria, per each pneumococcal strain) the area occupied by the green fluorescence signal of the bacteria or Ply was divided by the area occupied by the red fluorescence signal of β-actin. All areas were measured in square pixels and calculated with the software Image J. Columns in the graph represent average values, and error bars represent standard deviations. The Pneumococci/ β-actin ratio is shown on the Y axis; * = p<0.05, n.s. = non-significant.

To explore if anti-β-actin antibody treatment also affected neuronal cell death, treated and untreated neurons were infected with *S. pneumoniae* TIGR4 and stained with a live/dead dye. Through high-resolution live-cell imaging microscopy, we demonstrated that the neuronal cell death was significantly decreased after anti-β-actin antibody treatment (***Figure 8C***). To be closer to the human host setting, we also used primary neurons differentiated from human neuroepithelial stem cells. Immunofluorescence high-resolution microscopy revealed that β-actin is expressed also on the plasma membrane of primary neurons (***Figure 8D***) as previously reported (***Dinic et al., 2013***; ***Sechi et al, 2000***; ***Hoock et al., 1991***). Adherent TIGR4 bacteria were observed to co-localize with β-actin (***Figure 8D***). After one-hour treatment with anti-β-actin antibody, high-resolution microscopy and quantification analyses showed that the adherence of TIGR4 pneumococci to primary neurons was significantly impaired (***Figures 8D*** and ***8E***), while the mouse IgG isotype control treatment showed adherence levels similar to what was observed for untreated neurons (***Figures 8D*** and ***8E***). Adhesion of TIGR4Δ*rrgA* bacteria to primary neurons was significantly lower than for wt TIGR4, and the complemented strain TIGR4Δ*rrgA+rrgA* was not affected by β-actin blockade (***Figures 8D*** and ***8E***). Furthermore, there was no co-localization between TIGR4Δ*rrgA* bacteria and β-actin (***Figures 8D*** and ***8E***). Lastly, high-resolution microscopy and quantification analysis showed that anti-β-actin antibody treatment significantly reduced the interaction of Ply with the plasma membrane of neurons (***Figures 8F*** and ***8G***). Since the Ply signal can emanate from the cytosol of pneumococci, or from the bacterial surface or in a released form, often within extracellular vesicles (EVs) derived from pneumococci (***Codemo et al., 2018***), we cannot at this stage conclude whether the antibody blockade acts directly to prevent Ply binding to the plasma membrane of primary neurons or indirectly by preventing bacterial and/or EV-binding.

## Discussion

An acute episode of pneumococcal meningitis is frequently followed by a variety of neurological sequelae usually developing during the first 90 days after infection (***Lucas et al., 2016***; ***Schmidt et al., 2016***; ***Chandran et al., 2011***; ***Nau et al., 2002***). Symptoms may come from cranial nerve dysfunctions leading to hearing loss and vision impairment, or from motor neurons, leading to hemiplegia or ataxia. But neuronal sequelae may also result in epilepsy, and psychiatric disorders. The distinct types of neurological sequelae suggest that, in pneumococcal meningitis, bacteria may infect neuronal cells in the brain, leading to neuronal cell death, and that the types of sequelae reflect their localization in the brain and what function these missing neurons had (***Lucas et al., 2016***; ***Chandran et al., 2011***; ***Nau et al., 2002***).

In this study we demonstrate that the pneumococcal strain TIGR4 both adheres to and is taken up intracellularly by neurons leading to neuronal cell death. Adherence to neuronal cells was found to be primarily mediated by the pilus-1 adhesin RrgA. Our data further show that purified RrgA can interact with neuronal β-actin. The crystal structure of the pneumococcal pilus-1 associated RrgA adhesin has revealed that its D3-domain exhibits an integrin-like fold (***Izoré et al., 2010***). It is known that the actin cytoskeleton interacts with integrins of the extracellular matrix (ECM) and integrin-actin interaction is crucial for cellular adhesion to the ECM and cell-to-cell stability (***Vicente-Manzanares et al., 2009***). Furthermore, integrin adhesion complexes possess actin-polymerization activity (***Butler et al., 2006***). It is therefore here suggested that RrgA through domain D3 acts as an integrin mimic directly interacting with β-actin, allowing bacterial adhesion to membrane sites on neurons that can respond dynamically via actin polymerization. The direct interaction between domain D3 and β-actin remains though to be shown in future studies. The importance of RrgA for brain infection is also supported by previous studies demonstrating that RrgA promotes bacterial entry into the brain from the blood-stream by interacting with the two endothelial receptors PECAM-1 and pIgR, and at autopsy, after a fatal pneumococcal meningitis, five out of six brains studied contained RrgA expressing pneumococci (***Iovino et al, 2017***). However, only about 20-30% of clinical pneumococcal isolates harbour the pilus-1 islet (***Moschioni et al., 2010***; ***Iovino et al., 2020***). Thus, a larger study is required to be able to conclude whether expression of RrgA increases the risk for developing severe neurological sequelae following an episode of pneumococcal meningitis.

We found here that pneumococcal internalization by neurons requires both expression of RrgA and Ply. The cytotoxin Ply forms transmembrane pores at cholesterol rich sites in eukaryotic membranes, and recently it was demonstrated that domain 4 of Ply not only binds to cholesterol, but also to the mannose-receptor MRC-1, and that this interaction can lead to pneumococcal uptake by MRC-1 expressing dendritic cells and M2-polarized macrophages (***Subramanian et al., 2019***). We here provided evidence that Ply triggers neuronal uptake of pneumococci by directly interacting with the cytoskeleton protein β-actin. In neuronal cells, the actin cytoskeleton is connected to cholesterol rich lipid rafts and the dynamic interaction between lipid rafts and the actin cytoskeleton regulates many aspects of neuronal cells (***Head et al., 2014***). It is therefore here suggested that Ply can interact with β-actin in neuronal lipid rafts, leading to localized formation of cholesterol-dependent membrane pores, affecting the actin cytoskeletal dynamics at these sites, thereby promoting pneumococcal uptake through endocytosis.

We observed that pneumococci induce neuronal cell death that involves both bacterial adherence via RrgA and expression of Ply. The synergistic effects mediated by these two virulence factors suggest that bacterial uptake by neurons promotes further cell death. Ply is not actively secreted by pneumococci since it lacks a secretion signal sequence, unlike other pore-forming toxins. Instead the toxin is released through bacterial autolysis and as a cytosolic cargo in extracellular membrane vesicles frequently produced by pneumococci (***Codemo et al., 2018***).

The main bacterial localization of Ply is in the cytosol, but it was also recently found on the bacterial surface (***Subramanian et al., 2019***). We envision that pneumococcal entry from the circulation into the brain brings in bacterial associated Ply, and Ply already released in the blood stream. Bacterial associated cytotoxicity could explain why RrgA- and Ply-expressing pneumococci are considerably more neurotoxic than non-adhering Ply-expressing pneumococci, even though exogenously supplied Ply is neurotoxic even in the absence of bacteria (***Stringaris et al., 2002***; ***Braun et al., 2007***). Bacterial associated cytotoxicity might also contribute to localized uptake of pneumococci. Once internalised by neuronal cells, cytosolic Ply might be released through bacterial lysis, affecting internal neuronal membrane structures, potentially leading to increased influx of calcium from internal stores, similarly to what was previously observed for alpha-haemolysin in epithelial cells (***Uhlén et al., 2000***). RrgA-expressing pneumococci, not producing Ply, caused more neuronal cell death than bacteria lacking both virulence-factors. RrgA-expressing bacteria without Ply can still attach and become internalized and interact with neuronal β-actin leading to cell death. This suggests that bacterial binding to β-actin has cytotoxic effects, though to a lower degree than the cytotoxic effect of Ply. It is therefore possible that the pneumococcal RrgA-β-actin interaction perturbs the actin skeleton, thereby increasing the release of calcium from internal stores and/or activating other signal transduction pathways as we have previously demonstrated that purified RrgA stimulates macrophage motility through interaction with the CR3-integrin (***Orrskog et al., 2012***).

Furthermore, we found that pneumococcal expression of pilus-1 and mainly Ply enhances intracellular levels of calcium (Ca^2+^) in neurons. For eukaryotic cells, the interplay between the actin cytoskeleton and Ca^2+^ plays a fundamental role for cell growth and motility (***Bunnell et al., 2011***). Actin cytoskeleton proteins have been shown to inhibit Ca^2+^ mobilization from internal stores, a protective cellular response as high levels of Ca^2+^ contribute to neuronal cytotoxicity (***Gasperini et al., 2017***; ***Stueber et al., 2017***; ***Kass and Orrenius, 1999***). Purified Ply has been demonstrated to have neuro-toxic effects that involves calcium influx likely occurring via the pore-forming activity of the toxin (***Stringaris et al., 2002***). Streptolysin O (SLO), a cholesterol-dependent pore forming toxin like Ply, was shown to be removed from the plasma membrane by endocytosis in a membrane repair process requiring calcium (***Idone et al., 2008***). The increased calcium influx observed in TIGR4 infected neurons, depending on Ply expression, occurred concomitantly with an increased intracellular uptake of RrgA-expressing pneumococci. This suggest that localized membrane pore formation and calcium influx, in close proximity to β-actin to which the bacteria adhere, may activate local actin polymerization, resulting in endocytic uptake of pneumococci via a neuronal membrane repair process. This is supported by our finding that endocytosed pneumococci and Ply co-localize with β-actin. It is therefore possible that the pneumococcal-β-actin interaction perturbs the actin skeleton, thereby increasing the release of calcium from internal stores.

In conclusion, we have shown that pneumococci can interact with primary neurons as well as differentiated and non-differentiated neurons and induce neuronal death that is dependent on the presence of the pneumococcal pilus-1 adhesin RrgA and the cytotoxin Ply. Moreover, we found that RrgA promote binding to neurons, and together with Ply enhance pneumococcal entry into neurons. We also observed that both RrgA and Ply can interact with the cytoskeleton protein β-actin and that Ply increases intracellular calcium (Ca^2+^) levels in neurons. This suggests that RrgA and pneumolysin can promote disruption of β-actin filaments of neurons leading to cell death. Using anti-β-actin antibody we could inhibit pneumococcal uptake and cellular toxicity in the neurons. Since neurons usually do not regenerate, these data suggest that the two proteins and their interaction with β-actin could be important for why pneumococci may cause severe neurological sequelae. Further future studies are though required to confirm these findings in the human setting.

## Acknowledgments

We thank the iPS core facility at the Karolinska Institutet for isolation and differentiation of primary neurons, and the Mass Spectrometry Facility at Uppsala University for the mass spectrometry analysis. We thank Dr. Fariba Foroogh for the technical help with the cryostat tissue cutting, Dr. Peter Mellroth for providing us with the anti-GAPDH antiserum, Dr. Shigeaki Kanatani for the help provided during the Ca^2+^ imaging experiments, and Prof. Staffan Normark for the scientific discussions. We thank all the major funding that has supported this study: Petrus and Augusta Hedlund Foundation, Jeansson Foundation, Åke Wiberg Foundation, SFO StratNeuro Start-up grant, Clas Groschinsky Foundation and the Karolinska Institutet Research Grant Foundations, as well as the Swedish Research Council, the Knut and Alice Wallenberg Foundation, Stockholm County Council, and the Swedish foundation for Strategic research (SSF).

## Author Contributions

M.T. and F.I. designed the study; M.T., S.T., M.L., M.M. performed the experiments; all authors analyzed the data; M.T., B.H.N., and F.I. wrote the manuscript. All authors contributed to writing and have approved the final version of the manuscript.

## Material and Methods

### Culture of human cells

Neuroblastoma cells (SH-SY5Y, ATCC® CRL-2266™) were cultivated using Eagle’s Minimum Essential Medium (EMEM) and Ham’s F12 (Gibco) medium, 15% fetal bovine serum (FBS) (Thermo Fisher Scientific) and 1% Penicillin-Streptomycin (PIS) (Thermo Fisher Scientific) as previously described (Shipley et al., 2016). SH-SY5Y cells were differentiated using 10 µM Retinoic Acid (RA) (Bio-Techne) for 7 days, 50ng/ml Brain Derived Neuronal Factor (BDNF) (Sigma-Aldrich) was added at day 5 of differentiation (Shipley et al, 2016).

Primary neurons were obtained from the iPS Core Facility at Karolinska Institute. Briefly, neuroepithelial stem (NES) cells were seeded at low (20-30,000 cells/cm^2^) and high (40-50,000 cells/cm^2^) density in 24 well plates with glass coverslips, cultivated at 37°C/5% CO_2_ with DMEM/F-12+Glutamax (Gibco) and 1% PIS, induced with 1% N-2 (Gibco) and 0.1% B-27 (Gibco) supplement in every 2 days for 20 days for differentiation.

### Cultivation of *S. pneumoniae*

Strains used in the study include wild-type TIGR4, the isogenic deletion mutants of pilus-1 (TIGR4Δ*rrgA*-*srtD)* (Iovino et al., 2016b), RrgA (TIGR4Δ*rrgA)* (Iovino et al, 2016b) and pneumolysin (TIGR4Δ*ply)* (Subramanian et al., 2019). The double mutant strain T4Δ*rrgA-srt*DΔ*ply* was constructed by replacing the *ply* ORF with a kanamycin resistance gene in the previously constructed non-piliated strain T4Δ*rrgA-srtD* (Nelson et al., 2007). Briefly, the upstream region of the *ply* gene was amplified by polymerase chain reaction (PCR) using primers *ply-1* and *ply-2*, while the downstream region was amplified with primers *ply-3* and *ply-4*. The kanamycin resistance gene was obtained with primers *kanRfwd* and *kanRrev*. Primers *ply-2* and *ply-3* contain an overhang which overlaps with the *kanRfwd* and *kanRrev* primers, respectively. An overlap PCR reaction using primers *ply-1* and *ply-4* and the three DNA fragments yielded a single PCR product containing the *ply* region with a KanR ORF replacement which was then transformed into the T4Δ*rrgA-srt* strain and selected in blood agar plates with kanamycin (200 µg/mL). Deletion was confirmed via sequencing. Primers used are listed in Supplementary Table S4. All strains were plated on blood agar overnight at 37°C. Bacterial colonies were collected and grown in Todd-Hewitt broth (0.5% yeast extract) at 37°C and harvested at OD_600_=0.45-0.5, aliquots stored at −80°C as a 15% glycerol solution.

### Infection of neurons

SH-SY5Y cells were seeded at a density of 300,000 cells/well in 6 well plates for CFU adhesion assays; 900,000 cells/well in 6 well plates for CFU invasion assay. For Immunofluorescence, cells were seeded at a density of 150,000 cells/well in 12 well plates with a glass coverslip in seeding medium (1:1 EMEM: F12, 5% FBS). Incubated at 37°C/5% CO_2_ overnight, cells were then washed with PBS after overnight cultivation, replaced with new seeding medium and incubated for one hour prior to the infection assay.

Neuronal differentiation: SH-SY5Y cells were seeded at a density of 400,000 cells/well in 6 well plates without coverslip and 35,000 cells/well in 12 well plates with a glass coverslip in seeding medium (1:1 EMEM: F12, 5% FBS, 10µM RA). Incubated at 37°C/5% CO_2_ overnight, cells were washed with PBS after overnight cultivation, replaced with new seeding medium and incubated for one hour prior to the infection assay. Primary neurons were differentiated from NES cells in 24 wells with glass coverslips. At day 21 of differentiation, cells were washed once with PBS and replaced with seeding medium (DMEM/F-12, 5% FBS, 1%N-2, 0.1% B-27), then incubated for 1 hour at 37°C/5% CO_2_ prior to infection assay.

Adherence and uptake assays: SH-SY5Y cells, SH-SY5Y differentiated neurons and primary neurons were infected with multiplicity of infection (MOI) 10. Infected cells were incubated at 37°C /5% CO_2_ for two hours, and Colony Forming Unit (CFU) counting or Immunofluorescence staining was performed. CFU counting: To assess pneumococcal adhesion, after two hours of incubation, supernatant was collected from each well (non-adhered bacteria), wells were washed with PBS to eliminate unbound bacteria. Cells were then treated with 1 ml of Trypsin for 10∼15 min, cell suspensions were collected (adhered bacteria). Bacteria collected prior to and after adhesion were serial diluted in PBS and plated on blood agar at 37°C/5% CO_2_ overnight. To assess pneumococcal uptake into neurons, after two hours of incubation, wells were washed with PBS to eliminate unbound bacteria and treated with 150 μg Gentamycin (Gibco) in 1 ml of medium for one hour to kill extracellular bacteria. Cells were then washed with PBS and lysed with 1% Saponin (Sigma Aldrich) solution, cell suspensions were diluted with PBS and plated in blood agar at 37°C /5% CO_2_ overnight. Adhesion ratio was calculated as [CFU of adhered bacteria] / [CFU of (non−adhered bacteria + adhered bacteria)], uptake ratio was calculated as [CFU of internalized bacteria] / [CFU of adhered bacteria)]. Blockade of β-actin: differentiated neurons and primary neurons were treated with 2µg of either anti β-actin (mouse IgG1) Antibody (Thermo Fisher Scientific), or normal mouse IgG1(Abcam) as isotype control, or anti-ALK antibody (Thermo Fisher Scientific) as control for the specificity of β-actin blockade, centrifuged with slow rotation at 500 RPM for 5 minutes to enhance the binding of antibodies on the cell surface, then incubated for one hour at 37°C/5% CO_2_ prior to infection. Cells (differentiated neurons and primary neurons) were then infected with TIGR4 for 2 hours followed with CFU assay for differentiated neurons, and fixation followed by immunofluorescent staining for primary neurons.

### Mouse experiments

All animal experiments were approved by the local ethical committee (Stockholms Norra djurförsöksetiska nämnd). We used the bacteremia-derived meningitis model previously described by our group (Iovino et al., 2016a; Iovino et al., 2016b; Iovino et al., 2017; Iovino et al., 2018). Briefly, 5 male C57BL/6 wild-type mice 5 to 6 weeks old (Charles River) per experimental group were used that were anesthetized by inhalation of isofluorane (Abbott) before infection. 100 µl of 5×10^7^ CFU were injected intravenously into the tail and the mice were sacrificed at 10 hours post-infection. Clinical symptoms of the infected mice were observed during the infection according to ethical permit. After sacrifice, all mice were perfused to remove all bacteria still present in the blood of brain vessels, perfusion was performed as previously described (Iovino et al., 2016a; Iovino et al., 2016b; Iovino et al., 2017; Iovino et al., 2018). Brains were collected, cryopreserved in Shandon Cryomatrix (Thermo Fisher Scientific) and stored at −80°C.

### Immunofluorescence staining

After two hours of infection, human cells (SH-SY5Y, differentiated neurons, primary neurons) were washed with PBS then fixed with a 4% paraformaldehyde (PFA), permeabilized with 0.1% Triton X-100 in PBS and finally incubated with 1% Bovine serum albumin (BSA) solution in PBS for one hour. The permeabilization step was not performed for cell cultures used to detect bacterial adhesion on plasma membranes. Next, cells were incubated with primary antibodies for one hour and washed with PBS, followed by secondary antibodies for one hour in the dark and washed with PBS. Undifferentiated SH-SY5Y cells were stained with Phalloidin (Thermo Fisher Scientific) for one hour and washed with PBS, finally stained with DAPI for 10 minutes and washed with PBS. All antibodies were prepared using 2% BSA in PBS, 1:50 dilution for primaries, 1:200 for secondaries. Anti-capsule serotype 4 (rabbit) antibodies (Statens Serum Institute, Denmark) were used as primary antibodies for the detection of *S. pneumoniae* and followed with Alexa Fluor 488 goat anti rabbit antibody (Thermo Fisher Scientific). SH-SY5Y cells were stained with Phalloidin 594 (Thermo Fisher Scientific). Anti MAP2 (mouse IgG1) and gamma Enolase Antibodies (NSE-P1, mouse IgG1) (Santa Cruz Biotechnology) were used for staining of differentiated neurons and followed with Alexa Fluor 594 goat anti mouse antibody (Thermo Fisher Scientific). Anti β-actin (mouse IgG1) antibody (Thermo Fisher Scientific) used for staining of primary, differentiated neurons and mouse tissue, followed with Alexa Fluor 594 goat anti mouse antibody (Thermo Fisher Scientific). DAPI (Abcam) was used for all differentiated SH-SY5Y cells for staining process (1:5000). For imaging bacterial adhesion to SH-SY5Y, and differentiated neurons, a total of 500 cells with adhered bacteria were imaged per pneumococcal strain used. For imaging bacterial adhesion to primary neurons, a total of 100 cells with adhered bacteria were imaged per pneumococcal strain used. For imaging bacterial invasion of differentiated neurons, a total of 250 cells with intracellular bacteria were imaged. For imaging bacterial interaction with neurons in mouse brain sections, a total of 25 images per each section per each mouse (6 sections per each mouse, in total five mice per group) were taken.

*Ex vivo* mouse brain cryopreserved sections: frozen brains embedded in Cryomatrix (Thermo Fisher Scientific) were cut with a cryostat and 3 sections of 20 µm were placed on each microscope glass slide (VWR). Sections were fixed with acetone for 10 minutes, dried and incubated with anti-capsule serotype 4 antibody (rabbit IgG) mixed with anti-MAP2 antibody (mouse IgG) overnight at 4°C. Slides were washed with PBS and incubated for four hours with an Alexa Fluor 488 goat anti rabbit secondary antibody (for detection of *S. pneumoniae*) mixed with Alexa Fluor 647 goat anti mouse secondary antibody (for the detection of neuronal MAP2). Slides were washed with PBS and incubated for four hours with anti β-actin antibody labelled with 594 fluorophore using the Zenon Mouse IgG Labeling Kit (Thermo Fisher Scientific). Dilutions of primary and secondary antibodies were the same as the ones used for the staining of human cells described above. Vectashild (Vector Laboratories) was finally added to each coverslip with fixed cells or stained mouse brain section, covered with a coverslip and analyzed by fluorescence microscopy.

### High-resolution fluorescence microscopy and image processing

The high-resolution fluorescence microscope Delta Vision Elite Imaging System (GE healthcare) was used to analyze all immunofluorescence staining experiments. Images were acquired using a scientific complementary metal-oxide-semiconductor (sCMOS) camera and obtained using three different filters at different wavelengths: 488 nm (green), 594 nm (red) and 647 nm (purple, which we digitally changed into blue to have a better contrast with the green and red signals). Images were finally processed with Softworx imaging program (Applied Precision). For imaging adhered bacteria, human cells (SH-SY5Y, differentiated neurons and primary neurons) were fixed and not permeabilized, only the plasma membrane was imaged with a single snapshot for each filter. For imaging intracellular bacteria in differentiated neurons, or bacterial interaction with neurons in mouse brain sections, series of z-stacks were imaged in order to capture either the thickness of the neurons (in case of imaging differentiated neurons) or the cell morphology of neurons within each brain mouse section. The z-stacks Images (z-stacks) taken with the DV Elite Imaging System was rotated using the *3D Volume Viewer* function of the imaging software Softworx.

### Live-cell imaging experiments

Differentiated neurons were treated with 2μM Calcein AM and 4 μM Ethidium bromide from LIVE/DEAD™ Viability/Cytotoxicity Kit for 20 min at 37°C/5% CO_2_. Then cells were infected with different strains of TIGR4 at MOI=10 under a 20X objective microscope of the Delta Vision Elite Imaging System at 37°C/5% CO_2_, time lapse images were taken at every 10 seconds for total two hours with green, red and bright field channel.

### Ca^2+^ imaging of neurons

Ca^2+^ imaging of differentiated neurons was performed following the instructions of previously described set-up and manufacturer using Fluo-8 (AAT Bioquest) calcium indicator. Briefly, differentiated neurons were treated with 2μM Fluo-8 AM for 30 min at 37°C/5% CO2, then cells were washed once with medium and replaced with new medium without Fluo-8. Imaging of calcium was performed under a 20X objective microscope (Delta Vision Elite) using FITC fluorescence channel. Cells were infected with or without wt TIGR4 or its isogenic mutants at MOI=10 for two hours and then Fluo-8 AM was loaded for recording of calcium flux. In order to assess the effect of Ply on calcium flux, 100ng/mL of purified Ply was added to the cells (infected with TIGR4Δ*ply* for two hours and loaded with Fluo-8 for 30 min) 2 min before imaging. Image J was used for measurement of fluorescence intensity from manually chosen region of interest (ROI), intracellular Ca^2+^ levels were determined using ΔF/F_0_ as previously described (Uhlén et al., 2000). Originlab was used for background subtraction for all the calcium images.

### Quantification of fluorescent signal

The fluorescent signal intensity was quantified using Image J as previously described (Iovino et al., 2017). Briefly, the fluorescent signal to quantify was first converted into a grayscale image and inside each field the area covered by the fluorescent signal was selected using the function *Image-Adjust-Color Threshold*. The area covered by each fluorescent signal was then measured using the function *Analyze-Measure* (area measured in pixel^2^). For quantification of the fluorescent intensity (Figure 6B), the fluorescent signal was first converted into a grayscale image and using the function *Analyze-Plot Profile* of Image J the graph (pixel distance on the X axis, pixel intensity on the Y axis) was generated.

### Co-immunoprecipitation with RrgA, Ply and neuronal lysate or purified β-actin and α-tubulin 1B

Co-immunoprecipitations were performed using Ni-NTA magnetic beads (Thermo Fisher Scientific). First, Ni-NTA beads were equilibrated using a washing buffer containing 300 mM NaCl, 50 mM NaH2PO4, 20 mM imidazole, 1.5 mM MgCl2, 0.1% Triton×100, 3% glycerol and then incubated with 2 μM purified Ply, RrgA or RrgB for 30 min. For pull-down using neuronal cell lysate, 5×10^6^ differentiated neurons were lysed in lysis buffer containing 150 mM NaCl, 50 mM NaH2PO4, 10 mM imidazole, 1.5 mM MgCl2, 0.1% Triton×100, 0.6% of n-Dodecyl β-D-maltoside, 3% glycerol; 0.9 mM DTT, complete protease inhibitor EDTA-free (Sigma Aldrich), and homogenized by passing through a needle. The lysate was centrifuged at 3500 rpm for 10min at 4°C. The supernatant was incubated with empty or, Ply or RrgA-conjugated Ni-NTA beads at 4°C for 2 hrs. Beads were washed four times with washing buffer. The bound proteins were eluted by adding LDS sample buffer (Thermo Fisher Scientific) and incubating for 3 min at 55°C. Eluted proteins were loaded on SDS-PAGE for Coomassie Blue staining and sent for mass spectrometry analysis at the Science for Life Laboratory at Uppsala, Sweden. Proteins were identified according to the criteria of at least two matching peptides of 95% confidence level per protein. Western blot was done after the pull-down assay to confirm the binding of β-actin to RrgA and Ply. For Ni-NTA pull-down using purified proteins, 2 μM purified β-actin or α-tubulin 1B were incubated with empty or, Ply or RrgA-conjugated Ni-NTA beads at 4°C for 2 hrs. Beads were washed four times with washing buffer (containing 30mM instead 20mM imidazole). Washing steps performed using a buffer containing 300 mM NaCl, 50 mM NaH2PO4, 30 mM imidazole, 1.5 mM MgCl2, 0.1% Triton×100, 3% glycerol, The bound proteins were eluted by adding LDS sample buffer and incubating for 5 min at 95°C. Eluted proteins were loaded on SDS-PAGE for western blot analysis.

### Mass Spectrometry

Gel bands were cut into small (1 mm^3^ pieces), destained using acetonitrile (ACN), washed and exposed to dithiothreitol (DTT) reduction and iodoacetamide (IAA) alkylation. Thereafter the proteins were digested by sequencing grade modified trypsin at a concentration of 12.5 ng/µL in 25 mM ammonium bicarbonate pH 8 overnight at 37°C. The peptides were extracted by sonication in 60% ACN and 5% formic acid (FA). Finally, the extracted peptides were completely dried to completion and thereafter desalted using the SPE Pierce C18 Spin Columns (Thermo Fisher Scientific). These columns were activated by 2 × 200 μL of 50% acetonitrile (CAN) and equilibrated with 2 × 200 μL of 0.5% trifluoroacetic acid (TFA). The tryptic peptides were adsorbed to the media using two repeated cycles of 40 μL sample loading and the column was washed using 3 × 200 μL of 0.5% TFA. Finally, the peptides were eluted in 3 × 50 μL of 70% ACN and dried. Dried peptides were resolved in 40 μL of 0.1% formic acid and further diluted 4 times prior to nano-LC-MS/MS. The nanoLC-MS/MS experiments were performed using a Q Exactive Orbitrap mass spectrometer (Thermo Fisher Scientific) equipped with a nano electrospray ion source. The peptides were separated by C18 reversed phase liquid chromatography using an EASY-nLC 1000 system (Thermo Fisher Scientific). A set-up of pre-column and analytical column was used. The precolumn was a 2 cm EASYcolumn (ID 100 µm, 5 µm particles) (Thermo Fisher Scientific) while the analytical column was a 10 cm EASY-column (ID 75 µm, 3 µm particles, Thermo Fisher Scientific). Peptides were eluted with a 35 min linear gradient from 4% to 100% acetonitrile at 250 nL min-1. The mass spectrometer was operated in positive ion mode acquiring a survey mass spectrum with resolving power 70,000 (full width half maximum), m/z 400-1750 using an automatic gain control (AGC) target of 3×106. The 10 most intense ions were selected for higher-energy collisional dissociation (HCD) fragmentation (25% normalized collision energy) and MS/MS spectra were generated with an AGC target of 5×105 at a resolution of 17,500. The mass spectrometer worked in data-dependent mode. The acquired data RAW-files) were processed by Proteome Discoverer software (Thermo Scientific, version [nr 1.4.1.14]) using the Sequest algorithm towards a combined database containing protein sequences from Homo Sapience (26546 entries) downloaded from Uniprot 2019-06. For the identification of the neuronal proteins bound to RrgA and Ply, we have considered the proteins with a score higher than 50 (Supplementary Tables S1-3). All proteins with a score higher than 50 bound to RrgA and Ply were also present in the negative control. The protein in the negative control with the lowest score (score = 20.41, Supplementary Table S3) was β-actin which was present at much higher scores among the neuronal proteins bound to RrgA and Ply, with scores of 86.13 and 106.26 respectively (Supplementary Tables S1 and S2). The score of 20 is usually the threshold for false positive proteins (20), therefore in our case β-actin can be considered a false positive in the negative control highlighting the finding that β-actin could be a specific ligand of RrgA and Ply.

### Western blot analysis

3.5 × 10^6^ cells (differentiated neurons from SH-SY5Y cells) were harvested and lysed in a RIPA buffer containing protease and phosphatase inhibitors. Lysed cells were centrifuged for 10 min at 4°C, supernatants were collected and protein concentration was adjusted to 1μg/μl. Samples were boiled in 95°C for 10 minutes and loaded into NuPage Novex 4-12% Bis-Tris SDS-PAGE Gels (Thermo Fisher Scientific), electroblotting was performed using the Biorad Trans-Blot Turbo Transfer System. Membranes were first incubated overnight with PBS-T (T=0,1% Tween) supplemented with 5% milk. After washing the membrane with PBS-T, incubation with primary antibodies (1:2000) was performed for three hours, incubation with secondary antibodies (1:5000) was performed for one hour. Antibodies used for Western blot experiments were diluted in PBS-T supplemented with 1% dry milk. For data shown in Supplementary Figure S1B, a rabbit polyclonal anti-GAPDH antiserum was used for detection of GAPDH as loading control. Horseradish peroxidase conjugated Goat anti-Mouse IgG (GE Healthcare) diluted 1:5000 was used as secondary antibodies for the detection of Mouse IgGs.

### Statistical analysis

Statistical analyses were performed using Prism 5. For two-group comparisons, the non-parametric Wilcoxon’s rank sum test (also known as Mann-Whitney test) was used. For multiple comparisons (more than two groups), the nonparametric ANOVA test was used to assess the presence of the differences between the groups. The ANOVA test was then combined with the Dunn’s test to make pairwise comparisons.

