## Supplementary Figures 1-5, Supplementary Tables 1-4 for "Neuronal death in pneumococcal meningitis is triggered by pneumolysin and pilus-1 interactions with β-actin"

### Supplementary Data

**Supplementary Figure 1. Differentiation of SH-SY5Y cells into mature neurons**

Generation of mature differentiated neurons from human SH-SY5Y neuroblastoma cells was assessed by the expression of neuronal specific markers MAP2 and NSE via (**A**) immunofluorescence microscopy and (**B**) western blot analysis in which GAPDH was used as loading control. White scale bars in Figure S1A represent 100 µm, in the blot in Figure S1B the same protein concentration of both SH-SY5Y and neurons was loaded into the SDS-page gel. (**C**) Through phase contrast light microscopy analysis, we also observed that differentiated neurons displayed the neuronal typical cell-to-cell connections and axon formation (red arrows), black scale bars represent 100 µm.

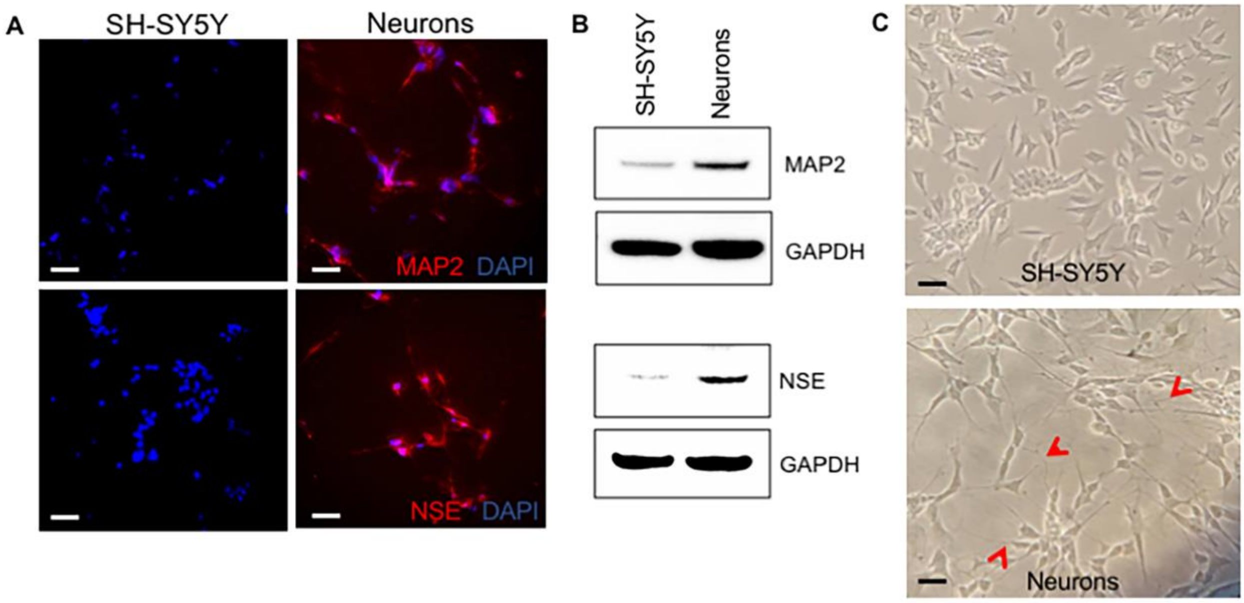

**Supplementary Figure 2. RrgA enhances pneumococcal adherence to SH-SY5Y cells, and RrgA and Ply increase pneumococcal uptake into SH-SY5Y cells.**

SH-SY5Y cells were challenged with pneumococci of MOI 10 and after 2 hours **(A)** adhesion and **(B)** uptake into the cells were measured. Strains used were wt TIGR4 and its isogenic mutants in the pilus, TIGR4Δ*rrgA-srtD*, *rrgA*, TIGR4Δ*rrgA*, and the *rrgA* mutant complemented with *rrgA*. TIGR4Δ*rrgA*+*rrgA*. (**C**) Adhesion and (D) uptake was measured using wt TIGR4 and its isogenic mutant in *ply*, TIGR4Δ*ply*. Adhesion ratio was calculated by dividing the total number of bacteria in each well for each pneumococcal strain after 2 hours infection by the total number of adhered bacteria in each well for each pneumococcal strain. For all graphs (A-D) the columns represent average values, and error bars represent standard deviations. Each graph shows data from at least three (n≥3) biological replicates. *** = p<0.0001, ** = p<0.001, ** = p<0.05, n.s.= not-significant.

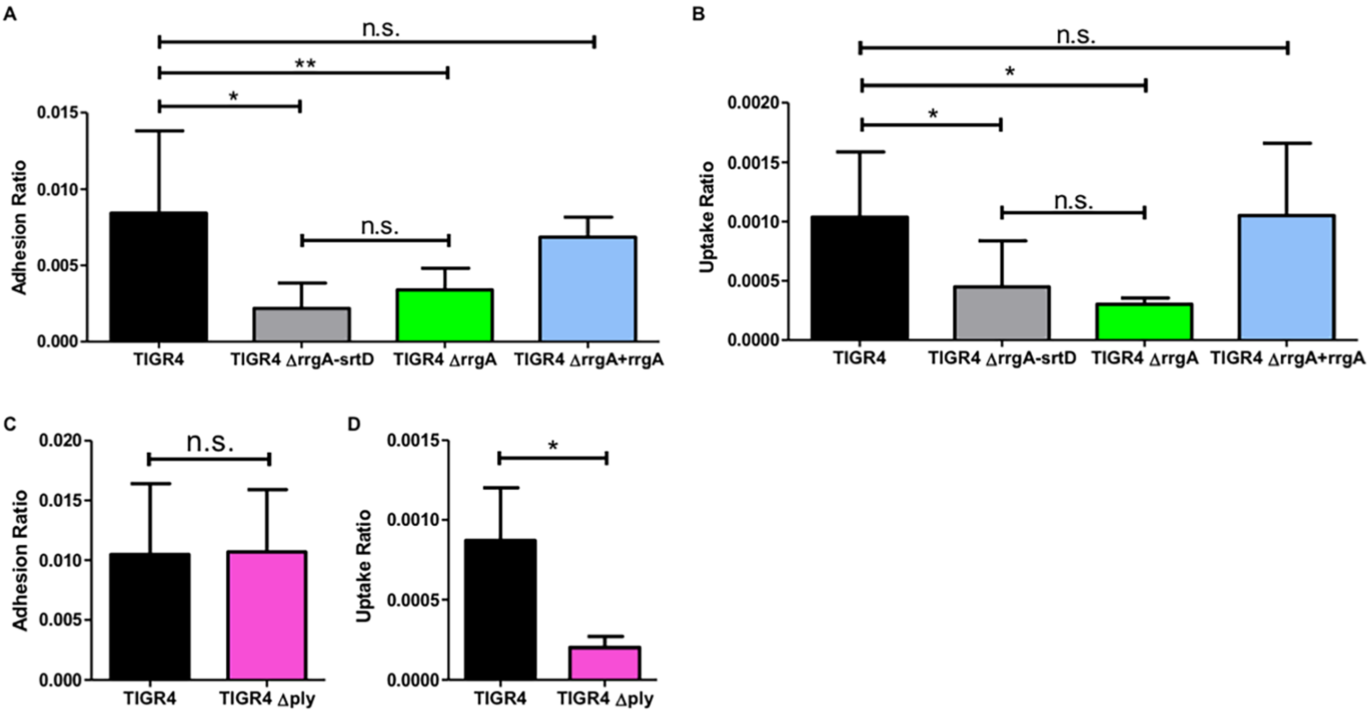

**Supplementary Figure 3. Imaging of piliated and non-piliated pneumococci that adhered to SH-SY5Y cells.**

(**A**) High-resolution fluorescence microscopy was used to visualize piliated and non-piliated pneumococci that adhered to SH-SY5Y cells. Neurons were stained with Phalloidin (red), and TIGR4 and TIGR4Δ*rrgA-srtD* were stained with anti-serotype 4 capsule antibody combined with goat anti rabbit Alexa Fluor 488 (green). White arrows point to pneumococci that adhered to SH-SY5Y cells. White scale bars represent 10 µm. The images shown are two representative images selected among 200 cells with adhered bacteria imaged per pneumococcal strain. The panel "Detail 5X" displays a 5X-magnified image of the area in the original images with bacteria that adhered to neurons. (**B**) Quantification of the number of bacteria that adhered to neurons based on the microscopy analysis results shown in Figure S1A. For quantification, the bacterial fluorescence signal on SH-SY5Y cells, in each image (n= 200 SH-SY5Y cells with adhered bacteria, per each pneumococcal strain) the area occupied by the green fluorescence signal of the bacteria, was divided by the area occupied by the red fluorescence signal of SH-SY5Y cells. All areas were measured in square pixels and calculated with the software Image J. The Pneumococci/Phalloidin ratio is shown on the Y axis. Columns in the graph represent average values, error bars represent standard deviations, * = p<0.05.

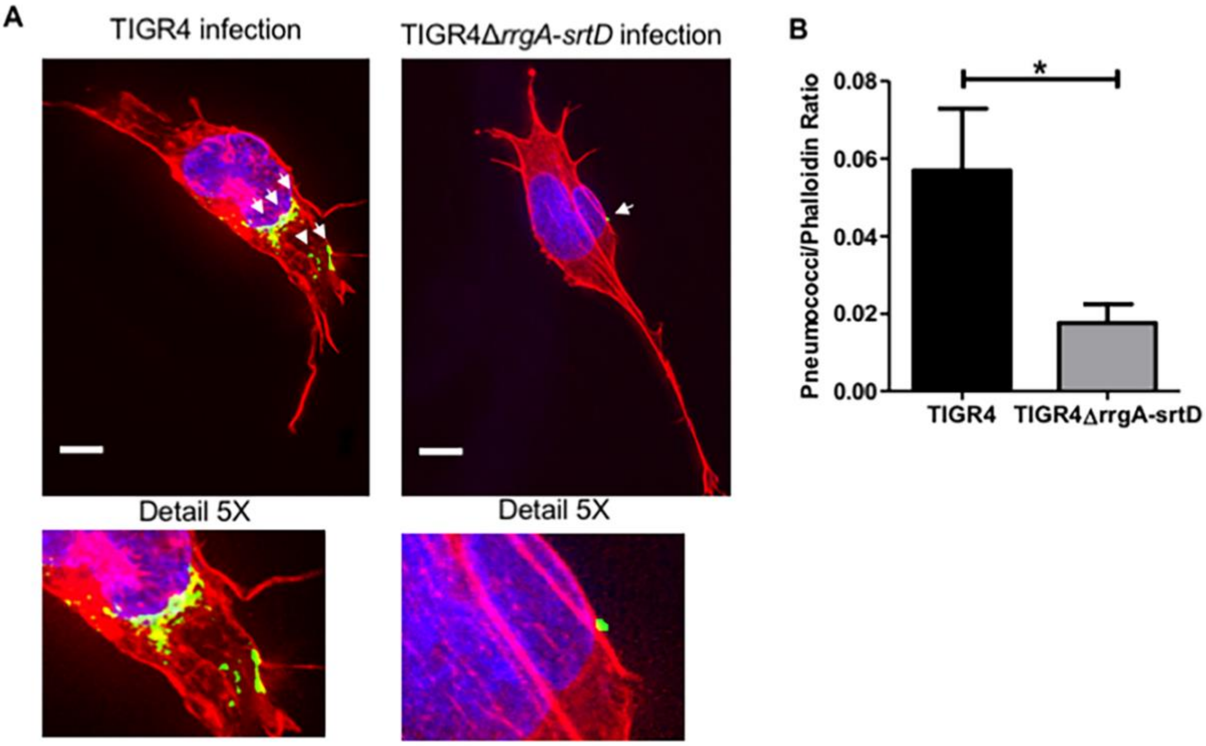

**Supplementary Figure 4. Coomassie staining of cell lysate of differentiated neurons**
Before performing the co-immunoprecipitation experiments, the quality of the cell lysate of differentiated neurons was assessed by SDS-page electrophoresis and Coomassie staining. The clear detection of the neuronal protein bands ranging from low to high molecular sizes suggested good quality of the neuronal cell lysate. The numbers on the left side of the image show the protein molecular weight in kDa.

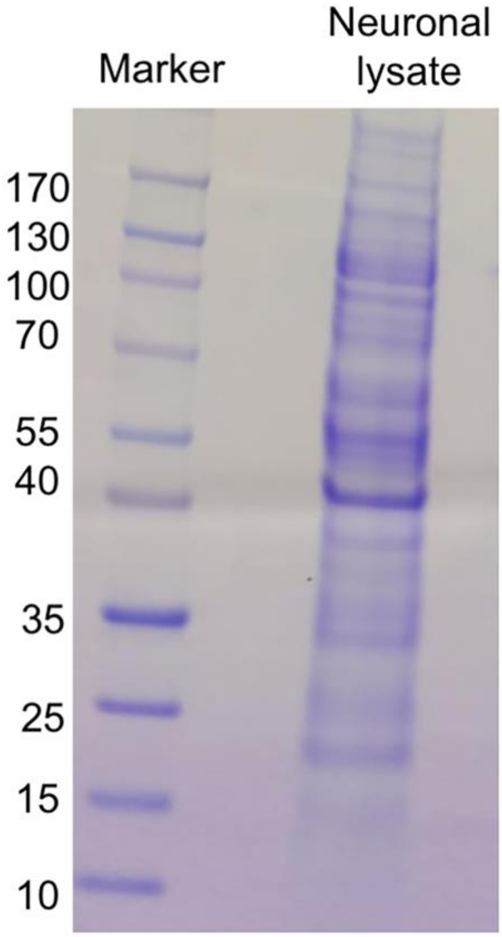

**Supplementary Figure 5. Neuronal cytoskeleton proteins identified by mass spectrometry** A pull-down assay was performed using differentiated neurons from SH-SY5Y cells and Ni-NTA beads-coupled-RrgA (RrgA), Ni-NTA beads-coupled-Ply (Ply) or Ni-NTA beads alone (Negative control) to identify proteins that bound to RrgA, or Ply respectively, as compared to beads alone. Mass spectrometry analysis identified the neuronal cytoskeleton proteins listed on the x axis and their presence is presented as Mass spectrometry scores on the Y axis. The dash black line at score = 20 represents the threshold of false positives.

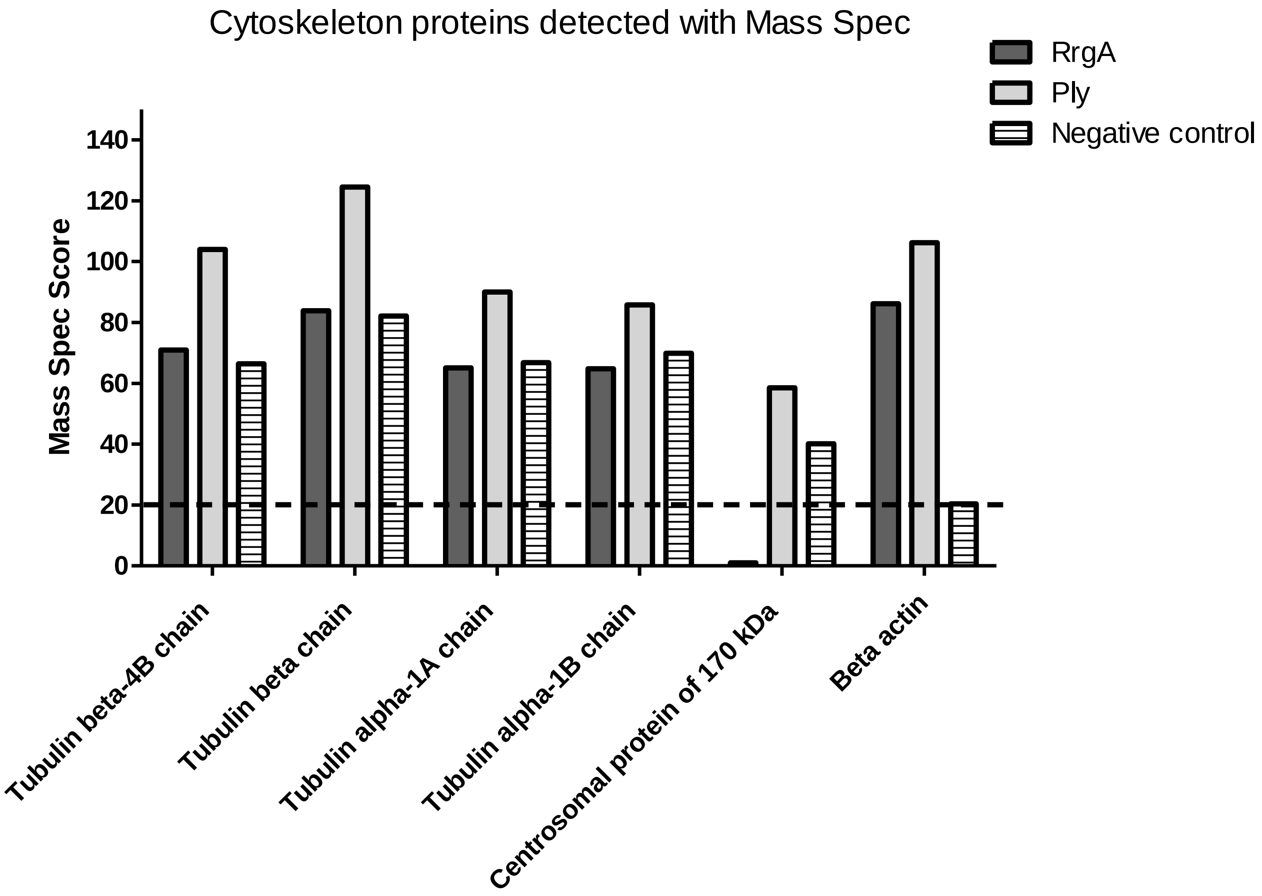

**Supplementary Table 1.** List of proteins bound to Ni-NTA-beads-coupled-RrgA identified by mass spectrometry. β-actin is shown in bold.

| **Neuronal proteins bound to RrgA** | **Score** | **Cellular localization** |
|---|---|---|
| Elongation factor Tu | 331.48 | Intracellular (mitochondria) |
| Non-POU domain-containing octamer-binding protein | 620.56 | Intracellular (nucleoplasm) |
| Splicing factor, proline- and glutamine-rich | 511.18 | Intracellular (DNA/RNA-binding) |
| Pre-mRNA-splicing factor ATP-dependent RNA helicase DHX15 | 207.21 | Intracellular (nuclear speckles) |
| Endoplasmic reticulum resident protein 44 | 118.42 | Intracellular (endoplasmatic reticulum) |
| Phospholipase D3 | 77.08 | Intracellular (lysosomes, endosomes, Golgi) |
| Cleavage and polyadenylation specificity factor subunit 7 | 86.06 | Intracellular (nucleoplasm) |
| 5'-nucleotidase domain-containing protein 2 | 111.90 | Intracellular (DNA binding) |
| Heterogeneous nuclear ribonucleoprotein H | 71.52 | Intracellular (nucleoplasm) |
| L-lactate dehydrogenase B chain | 98.82 | Intracellular (cytosol) |
| C-terminal-binding protein 1 | 54.12 | Intracellular (nucleoplasm) |
| DnaJ homolog subfamily C member 10 | 85.61 | Intracellular (chaperone, mitochondria) |
| Heterogeneous nuclear ribonucleoprotein L-like | 72.87 | Intracellular (nucleus) |
| Cleavage and polyadenylation specificity factor subunit 5 | 59.16 | Intracellular (RNA binding) |
| Paraspeckle component 1 | 72.24 | Intracellular (RNA binding) |
| Heterogeneous nuclear ribonucleoprotein H2 | 52.18 | Intracellular (RNA binding) |
| Aflatoxin B1 aldehyde reductase member 2 | 64.57 | Intracellular (Golgi) |
| Thioredoxin-dependent peroxide reductase, mitochondrial | 51.53 | Intracellular (mitochondria) |
| Cleavage and polyadenylation specificity factor subunit 6 | 52.68 | Intracellular (RNA binding) |
| Neurosecretory protein VGF | 129.68 | Secreted |
| Cytosolic purine 5'-nucleotidase | 50.04 | Intracellular (cytosol) |
| Putative RNA-binding protein Luc7-like 2 | 58.74 | Intracellular (RNA binding) |
| Tubulin beta-4B chain | 70.96 | Cytoskeleton |
| Prelamin-A/C | 51.3 | Intracellular (nucleus) |
| Tubulin beta chain | 83.9 | Cytoskeleton |
| Glutamate dehydrogenase 1, mitochondrial | 62.06 | Intracellular (mitochondria) |
| Tubulin alpha-1A chain | 65.09 | Cytoskeleton |
| Dihydropyrimidinase-related protein 4 | 60.43 | Intracellular (cytoplasm) |
| Heat shock cognate 71 kDa protein | 60.08 | Intracellular (nucleus) |
| Secretogranin-1 | 71.2 | Secreted |
| Dihydropyrimidinase-related protein 5 | 80.03 | Intracellular (cytoplasm) |
| Tubulin alpha-1B chain | 64.82 | Cytoskeleton |
| Splicing factor 1 | 51.24 | Intracellular (nucleus) |
| L-lactate dehydrogenase A chain | 62.3 | Intracellular (cytoplasm) |
| ATP-dependent RNA helicase DDX42 | 65.93 | Intracellular (RNA binding) |
| Heterogeneous nuclear ribonucleoprotein L | 125.64 | Intracellular (nucleus) |
| **Beta-actin** | **86.13** | **Cytoskeleton** |

**Supplementary Table 2.** List of proteins bound to Ni-NTA beads-coupled-Ply identified by mass spectrometry. β-actin is shown in bold.

| **Neuronal proteins bound to Ply** | **Score** | **Cellular localization** |
|---|---|---|
| Elongation factor Tu | 355.91 | Intracellular (mitochondria) |
| Non-POU domain-containing octamer-binding protein | 778.22 | Intracellular (nucleoplasm) |
| Splicing factor, proline- and glutamine-rich | 650.26 | Intracellular (DNA/RNA-binding) |
| Pre-mRNA-splicing factor ATP-dependent RNA helicase DHX15 | 201.65 | Intracellular (nuclear speckles) |
| Endoplasmic reticulum resident protein 44 | 125.30 | Intracellular (endoplasmatic reticulum) |
| Cleavage and polyadenylation specificity factor subunit 7 | 90.01 | Intracellular (nucleoplasm) |
| Heterogeneous nuclear ribonucleoprotein H | 79.37 | Intracellular (nucleoplasm) |
| Neurosecretory protein VGF | 164.31 | Secreted |
| ATP-dependent 6-phosphofructokinase, muscle type | 84.26 | Intracellular (cytosol, nucleus) |
| Heterogeneous nuclear ribonucleoprotein L-like | 86.22 | Intracellular (nucleoplasm) |
| Creatine kinase U-type, mitochondrial | 55.82 | Intracellular (mitochondria) |
| Glutamate dehydrogenase 1, mitochondrial | 91.65 | Intracellular (mitochondria) |
| Secretogranin-1 | 101.75 | Intracellular (endoplasmatic reticulum) |
| Cleavage and polyadenylation specificity factor subunit 5 | 65.97 | Intracellular (RNA binding) |
| Paraspeckle component 1 | 65.22 | Intracellular (RNA binding) |
| Aflatoxin B1 aldehyde reductase member 2 | 69.72 | Intracellular (Golgi) |
| Cleavage and polyadenylation specificity factor subunit 6 | 59.87 | Intracellular (RNA binding) |
| Cytosolic purine 5'-nucleotidase | 66.97 | Intracellular (cytosol) |
| Putative RNA-binding protein Luc7-like 2 | 61.14 | Intracellular (RNA binding) |
| Tubulin beta-4B chain | 103.96 | Cytoskeleton |
| Prelamin-A/C | 58.51 | Intracellular (nucleus) |
| Tubulin beta chain | 124.54 | Cytoskeleton |
| Tubulin alpha-1A chain | 90.04 | Cytoskeleton |
| Dihydropyrimidinase-related protein 4 | 76.7 | Intracellular (cytoplasm) |
| Dihydropyrimidinase-related protein 5 | 100.75 | Intracellular (cytoplasm) |
| Tubulin alpha-1B chain | 85.79 | Cytoskeleton |
| Splicing factor 1 | 62.5 | Intracellular (nucleus) |
| L-lactate dehydrogenase A chain | 69.29 | Intracellular (cytoplasm) |
| ATP-dependent RNA helicase DDX42 | 79.04 | Intracellular (RNA binding) |
| Heterogeneous nuclear ribonucleoprotein L | 86.22 | Intracellular (nucleus) |
| Zinc finger CCCH-type antiviral protein 1-like | 60.22 | Intracellular (cytoplasm) |
| Cystathionine beta-synthase | 63.36 | Intracellular (nucleus, cytoplasm) |
| Centrosomal protein of 170 kDa | 58.51 | Cytoskeleton |
| 5'-nucleotidase domain-containing protein 2 | 68.43 | Intracellular (cytplasm, nucleus, endoplasmatic reticulum) |
| Heat shock cognate 71 kDa protein | 53.9 | Intracellular (nucleus) |
| L-lactate dehydrogenase B chain | 56.22 | Intracellular (cytoplasm) |
| **Beta-actin** | **106.26** | **Cytoskeleton** |

**Supplementary Table 3.** List of proteins bound un-specifically to Ni-NTA beads identified by mass spectrometry. β-actin is shown in bold.

| **Negative control** | **Score** | **Cellular localization** |
|---|---|---|
| Elongation factor Tu | 355.91 | Intracellular (mitochondria) |
| Non-POU domain-containing octamer-binding protein | 778.22 | Intracellular (nucleoplasm) |
| Splicing factor, proline- and glutamine-rich | 439.83 | Intracellular (DNA/RNA-binding) |
| Pre-mRNA-splicing factor ATP-dependent RNA helicase DHX15 | 152.14 | Intracellular (nuclear speckles) |
| Endoplasmic reticulum resident protein 44 | 78.22 | Intracellular (endoplasmatic reticulum) |
| Phospholipase D3 | 37.31 | Intracellular (lysosomes, endosomes, Golgi) |
| Cleavage and polyadenylation specificity factor subunit 7 | 47.70 | Intracellular (nucleoplasm) |
| 5'-nucleotidase domain-containing protein 2 | 74.24 | Intracellular (DNA binding) |
| Heterogeneous nuclear ribonucleoprotein H | 53.09 | Intracellular (nucleoplasm) |
| L-lactate dehydrogenase B chain | 71.13 | Intracellular (cytosol) |
| Neurosecretory protein VGF | 121.05 | Secreted |
| ATP-dependent 6-phosphofructokinase, muscle type | 50.82 | Intracellular (cytosol, nucleus) |
| Heterogeneous nuclear ribonucleoprotein L-like | 53.09 | Intracellular (nucleoplasm) |
| Creatine kinase U-type, mitochondrial | 25.40 | Intracellular (mitochondria) |
| Glutamate dehydrogenase 1, mitochondrial | 62.22 | Intracellular (mitochondria) |
| Secretogranin-1 | 74.12 | Intracellular (endoplasmatic reticulum) |
| Cleavage and polyadenylation specificity factor subunit 5 | 40.59 | Intracellular (RNA binding) |
| Paraspeckle component 1 | 56.10 | Intracellular (RNA binding) |
| Heterogeneous nuclear ribonucleoprotein H2 | 37.30 | Intracellular (RNA binding) |
| Aflatoxin B1 aldehyde reductase member 2 | 52.99 | Intracellular (Golgi) |
| Thioredoxin-dependent peroxide reductase, mitochondrial | 41.85 | Intracellular (mitochondria) |
| Cleavage and polyadenylation specificity factor subunit 6 | 43.58 | Intracellular (RNA binding) |
| Cytosolic purine 5'-nucleotidase | 43.37 | Intracellular (cytosol) |
| Putative RNA-binding protein Luc7-like 2 | 52.84 | Intracellular (RNA binding) |
| Tubulin beta-4B chain | 66.38 | Cytoskeleton |
| Prelamin-A/C | 49.16 | Intracellular (nucleus) |
| Tubulin beta chain | 82.12 | Cytoskeleton |
| Glutamate dehydrogenase 1, mitochondrial | 62.22 | Intracellular (mitochondria) |
| Tubulin alpha-1A chain | 66.82 | Cytoskeleton |
| Dihydropyrimidinase-related protein 4 | 84.8 | Intracellular (cytoplasm) |
| Heat shock cognate 71 kDa protein | 62.49 | Intracellular (nucleus) |
| Dihydropyrimidinase-related protein 5 | 84.8 | Intracellular (cytoplasm) |
| Tubulin alpha-1B chain | 69.88 | Cytoskeleton |
| Splicing factor 1 | 57.59 | Intracellular (nucleus) |
| L-lactate dehydrogenase A chain | 72.1 | Intracellular (cytoplasm) |
| ATP-dependent RNA helicase DDX42 | 79.82 | Intracellular (RNA binding) |
| Heterogeneous nuclear ribonucleoprotein L | 147.03 | Intracellular (nucleus) |
| Zinc finger CCCH-type antiviral protein 1-like | 35.79 | Intracellular (cytoplasm) |
| Cystathionine beta-synthase | 39.93 | Intracellular (nucleus, cytoplasm) |

| | | |
|---|---|---|
| Centrosomal protein of 170 kDa | 40.14 | Cytoskeleton |
| **Beta-actin** | **20.41** | **Cytoskeleton** |

**Supplementary Table 4.** Primers used in this study.

| Primer Name | Sequence (5' – 3') |
|---|---|
| *ply-1* | GCTACCTGTCGCCCTTGCTC |
| *ply-2* | GATATTCTCATTTTAGCCATCTTCTACCTCCTAATAAGTTC |
| *ply-3* | ACTGGATGAATTGTTTTAGGAGAGGAGAATGCTTGCGAC |
| *ply-4* | GCTTGTTTAGCACGGTCGATAAC |
| *kanRfwd* | ATGGCTAAAATGAGAATATC |
| *kanRrev* | CTAAAACAATTCATCCAGT |
